# Response inhibition in premotor cortex corresponds to a complex reshuffle of the mesoscopic information network

**DOI:** 10.1101/2021.03.15.435381

**Authors:** Giampiero Bardella, Valentina Giuffrida, Franco Giarrocco, Emiliano Brunamonti, Pierpaolo Pani, Stefano Ferraina

**Affiliations:** Department of Physiology and Pharmacology, Sapienza University of Rome, 00185, Rome, Italy; PhD Program in Behavioral Neuroscience, Sapienza University of Rome, 00185, Rome, Italy

**Keywords:** information theory, motor control, inhibition, brain networks, complexity, behavior

## Abstract

Recent studies have explored functional and effective neural networks in animal models; however, the dynamics of information propagation among functional modules under cognitive control remain largely unknown. Here, we addressed the issue using Transfer Entropy and graph theory methods on mesoscopic neural activities recorded in the dorsal premotor cortex of rhesus monkeys. We focused our study on the decision time of a Stop-signal task, looking for patterns in the network configuration that could influence motor plan maturation when the Stop signal is provided. When comparing trials with successful inhibition to those with generated movement, the nodes of the network resulted organized into four clusters, hierarchically arranged, and distinctly involved in information transfer. Interestingly, the hierarchies and the strength of information transmission between clusters varied throughout the task, distinguishing between generated movements and canceled ones and corresponding to measurable levels of network complexity. Our results suggest a putative mechanism for motor inhibition in premotor cortex: a topological reshuffle of the information exchanged among ensembles of neurons.

**AUTHOR SUMMARY:** In this study, we investigated the dynamics of information transfer among functionally identified neural modules during cognitive motor control. Our focus was on mesoscopic neural activities in the dorsal premotor cortex of rhesus monkeys engaged in a Stop-signal task. Leveraging multivariate Transfer Entropy and graph theory, we uncovered insights on how behavioral control shapes the topology of information transmission in a local brain network. Task phases modulated the strength and hierarchy of information exchange between modules, revealing the nuanced interplay between neural populations during generated and canceled movements. Notably, during successful inhibition, the network displayed a distinctive configuration, unveiling a novel mechanism for motor inhibition in the premotor cortex: a topological reshuffle of information among neuronal ensembles.

## INTRODUCTION

The brain is a complex system with billions of neurons interacting across various spatial and temporal scales, forming interconnected modules of different sizes. At the small spatial scale, modules correspond to single neurons; at the large scale, modules correspond to specialized brain areas formed by many neurons. At the mesoscale, modules are neural populations of variable sizes, sometimes described as functional columns. Throughout the recent decades, neuroscience has attempted to characterize brain computations by relating neural activities to behavior. Understanding how network modules interact at the different scales and how information is shared and processed among parts is critical at all levels of investigation. Historically, most approaches have relied on single unit activity to describe the neural correlates of behavior in discrete populations of neurons recorded in a single area (Caminiti, Johnson, Galli, Ferraina, & Burnod, 1991; Fetz, 1992; Georgopoulos, Kalaska, Caminiti, & Massey, 1982; Munoz & Wurtz, 1993; Riehle, Grün, Diesmann, & Aertsen, 1997; Vaadia, Kurata, & Wise, 1988) being able to characterize larger networks only occasionally(Ferraina, Paré, & Wurtz, 2002; Wurtz, Sommer, Paré, & Ferraina, 2001). Recent developments in microelectrode fabrication (Hong & Lieber, 2019) have made it possible to record simultaneously from multiple channels and then investigate the dynamics of massive numbers of neurons and sites (Shenoy, Sahani, & Churchland, 2013). Nonetheless, the majority of studies focused on single-unit activity (**SUA**), and on the lower bands of the local field potentials (**LFPs**), which, however, indicate more the average synaptic input to the investigated area than the local spiking activity (for a review see (Herreras, 2016)). Comparatively, few studies have looked at signals that sample the average spiking activity of the population of neurons in close proximity to the recording electrode (multiunit activity; **MUA** (Mattia et al., 2013; Stark & Abeles, 2007; Supèr & Roelfsema, 2005; Trautmann et al., 2019)). Such mesoscopic signals, more stable than SUA, can provide crucial information for investigating the local dynamics that enable brain function control.

In the arm motor domain, studies have indicated that neurons in the dorsal premotor (PMd) cortex of monkeys are engaged in motor control, as evidenced by modulation patterns of both SUA and mesoscopic signals (LFPs and MUA) obtained both at the single and population level (Cisek & Kalaska, 2010; Milekovic, Truccolo, Grün, Riehle, & Brochier, 2015; Shenoy et al., 2013; Wise, Boussaoud, Johnson, & Caminiti, 1997). Some studies have employed the Stop signal task (Logan & Cowan, 1984), that requires actively inhibiting an upcoming movement upon presentation of a Stop signal, to investigate how neuronal activity in this area supports both movement generation and cancellation. According to these studies, neuronal activity in PMd is strongly related to both the generation and the suppression of an arm movement at different scales, from single neurons (Marcos et al., 2013; Mirabella, Pani, & Ferraina, 2011) to population of neurons(Pani et al., 2022) and local mesoscopic signals (Bardella, Pani, Brunamonti, Giarrocco, & Ferraina, 2020; Mattia et al., 2013; Pani et al., 2014). One of the proposed mechanisms suggests that arm movements are generated when neurons, following a Go signal, accumulate evidence up to a threshold, and inhibited when the accumulation is disrupted or postponed by a competing process triggered if a Stop signal is presented early enough after the Go signal. This view, in which neurons work as stochastic accumulators, stemmed from the analogy with the neural results in the oculomotor domain (Hanes, Patterson, & Schall, 1998; Paré & Hanes, 2003) and the original behavioral model (Logan & Cowan, 1984) based on a competition between accumulating processes. Recently (Pani et al., 2022), it has been advanced the hypothesis that such evolution can be depicted as a dynamic in the state-space, in which movement generation and inhibition correspond to distinct neural trajectories hopping within specific subspaces. While these approaches have been fruitful in giving a mechanistic (or quasi-mechanistic) account of movement control, the issue can be tackled also from another perspective. For example, an alternative hypothesis is that movement generation necessitates the functional interactions of clusters of neurons to build networks that efficiently handle information locally and send information to other brain areas and/or the spinal cord (see Bassett & Sporns review (Bassett & Sporns, 2017)). Thus, one could also ask whether motor decisions are characterized by specific routings of information across mesoscale neuronal “components”. To answer, we studied the PMd’s network dynamics using the MUA as a mesoscopic measure of spiking activity and implementing a combined information-theory and graph-theory approach to describe how the collective interaction of recorded modules is linked to motor decision-making. We demonstrated that in the behavioral epoch when decisions are made, mesoscopic modules may be classified into four distinct clusters that interact with one another and are organized hierarchically. Interestingly, the hierarchical level of some of them, as well as their strength of interaction and the overall communication rules, exhibited differences when movements are normally executed and when the animal successfully canceled the planned movement. Our results indicate that neurons in PMd have diverse opportunities to contribute to motor control at different spatial scales. In general, these findings support the idea of complexity being a core feature of the brain’s ability in managing information at multiple levels of neural interaction.

## MATERIALS AND METHODS

### Subjects

The study used two male rhesus macaque monkeys (Macaca mulatta; Monkeys P and C), weighing 9 and 9.5 kg, respectively. The animals’ care, housing, surgical operations, and experiments were all in accordance with European (Directive 86/609/ECC and 2010/63/UE) and Italian (D.L. 116/92 and D.L. 26/2014) laws and were approved by the Italian Ministry of Health. The monkeys were housed in pairs and given cage enrichment. During the recording days, they were fed regular primate chow supplemented with nuts and fresh fruits as needed, and they were given a daily water supply.

### Apparatus and task

The monkeys were placed in a soundproof, dimly lighted chamber in front of an isoluminant black background with a brightness intensity of less than 0.1 cd/m2. They were shown stimuli on a 17-inch touchscreen panel with an LCD display with an 800 × 600 pixel resolution. CORTEX, a non-commercial software program available at http://www.nimh.gov.it, was used to control the presentation of stimuli and recording of behavioral responses. Figure 1 depicts the overall task, which our group first described as a stop-signal reaching task (Mirabella et al., 2011). The central target (CT), a red circle with a diameter of 1.9 cm, was displayed at the start of each trial. The CT had to be reached by the monkeys to initiate the trial. A peripheral target (PT) randomly appeared on the left or right side of the screen after a variable holding duration of 400–900 ms, in steps of 100 ms. At the end of the holding period the CT vanished, signaling a Go signal. In the Go trials, the monkeys had to reach and hold on the PT for a range of 400–800 ms, in steps of 100 ms, to get a juice reward. The time between the presentation of the Go signal and the beginning of the hand movement was used to compute the reaction time (RT). The sequence of events in Stop trials was the same as in Go trials until the Go signal. The CT reappeared as the Stop signal after a variable delay known as the Stop signal delay (SSD), and the monkeys had to hold the CT until the end of the trial, which was between 800-1000 ms, to get the reward for a correct Stop trial. If the monkey removed its hand before the trial ended, it was considered an incorrect response for a Stop trial. For both correct Stop and correct Go trials, the same amount of juice was provided. The inter-trial interval was set at 800 milliseconds. Stop trials accounted for approximately 25% of all Monkey P trials and 32% of Monkey C trials. White rings around the central target provided input to the animals for the touch. A staircase tracking approach was utilized to determine the duration of the SSDs, to obtain a success rate of roughly 50% in Stop trials. The SSD rose by 100 ms in the succeeding Stop trial if the monkey successfully withheld the movement, but it reduced by 100 ms if the monkey failed to cancel the response.

**Figure 1.**
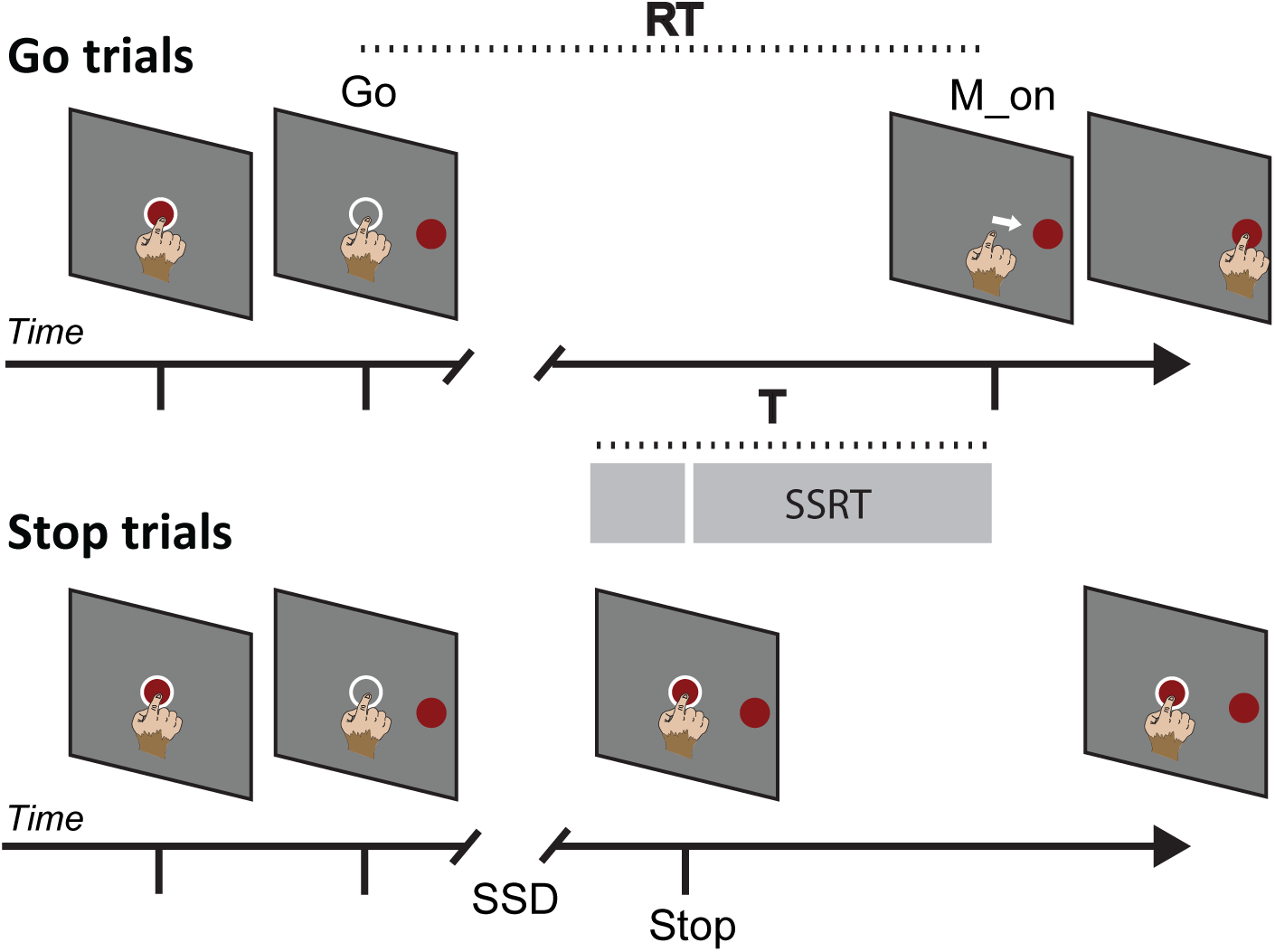
Sequence of Behavioral events characterizing the task. Go and Stop trials were randomly intermixed during each session. The epoch (**T**) of analysis is shown as a gray bar for both trials. White circles are feedbacks for the touch provided to the animals. **RT**, reaction time; **SSD**, Stop signal delay; **SSRT**, Stop signal reaction time; **Go**: time of Go signal appearance; **Stop**: Stop Signal appearance; **M ON**: movement onset.

### Behavioral considerations

The stop-signal reaching task allows for the calculation of the Stop signal reaction time (SSRT), which is recognised as a metric of efficiency in movement suppression. The race model(Logan & Cowan, 1984) is used to describe the behavior in the Stop trials, which involves two stochastic processes racing towards a threshold: the GO process, which is triggered by the Go signal and whose duration is represented by the RT, and the STOP process, whose duration must be derived. If the GO process wins the race against the STOP process, the movement is generated (resulting in wrong Stop trials). If instead the STOP process prevails, the movement is restrained (resulting in correct Stop trials). We may estimate the SSRT using the race model by considering the duration of the GO process, the probability of response, and the stop-signal delays (SSDs). To estimate the SSRT, we used the integration approach, which has been demonstrated to be the most trustworthy (Band, van der Molen, & Logan, 2003). This technique takes into account the probability of response [p(response)] for a given SSD in Stop trials. The longer the SSD, the higher the p(response), indicating a higher likelihood of a wrong Stop trial. By integrating p(response) over the distribution of Go trials, we can derive the nth value Go RT. Subtraction of the SSD from the nth Go RT yields the SSRT. To visualize the procedure, imagine an SSD where p (response) is exactly 0.5. For such SSD, the Go and Stop process would have the same probability of winning the race, meaning that the Stop and Go processes will both reach the threshold at the same time (from a probabilistic point of view). The completion times of the Stop and Go processes are, respectively, the end of the SSRT and the end of the RT (i.e., the beginning of movement). In this study, we compared the evolution of brain activity during movement preparation in the Go trials to the inhibitory effect of the Stop signal in correct Stop trials. To make this comparison effective, we considered an epoch before movement generation in Go trials and an epoch starting before the Stop signal and ending at the conclusion of the Stop process (SSRT duration) in correct Stop trials (see below). When adding wrong Stop trials in the analysis, we used the same epoch duration and we aligned neural activities to the beginning of movement. Because different SSDs were used in separate recording sessions, and because RT distributions also varied, resulting in varying SSRTs across sessions, we chose an epoch that could contain all estimated SSRTs in the various sessions (see Table S1). We decided for a duration of 400 ms for the epoch of analysis (Fig.1, epoch T, gray bar). In Stop trials, the epoch T did not include the Go signal when short SSDs were employed.

### Extraction and processing of neuronal data

In the left PMd, a multielectrode array (Blackrock Microsystems, Salt Lake City) with 96 electrodes spaced at 0.4 mm was surgically implanted. After opening the dura, the arcuate sulcus and precentral dimple were employed as reference (for electrode location see Figure S1 in (Pani et al., 2022)). We recorded the raw signal using a Tucker Davis Technologies (Alachua, FL) system, sampling at 24.4 kHz. As a measure of neuronal activity at the population level, local spectral estimated MUA was extracted offline from the raw signal. This was accomplished by computing the time-varying power spectra P(*ω*, t) from the signal’s short-time Fourier transform in 5-ms sliding windows, as described in Mattia et al. (Mattia et al., 2013). Relative spectra R(*ω*, t) were obtained normalizing P(*ω*, t) by their average P_ref_ (*ω*) across a fixed window (30 minutes) for the entire recording. Thus, the average R(*ω*, t) across the *ω*/2*π* band [0.2, 1.5] *kHz* represents our mesoscopic measure of spiking activity. As explained in Mattia et al. (Mattia et al., 2013), such estimation relies on two hypotheses. The first is that high *ω* components of the raw signal result from the convolution of firing rates *ν*(t) of neurons that are close to the electrode tip with a stereotypical single-unit waveform. The Fourier transform K(*ω*) of such an unknown waveform is canceled out in R(*ω*, t), which is therefore a good approximation of the ratio of firing rate spectra *|ν*(*ω*,t)*|*^2^ /*|ν*_ref_(*ω*)*|*^2^. The second is that high *ω* power *|ν*(*ω*,t)*|*^2^ is proportional to the firing rate *ν*(t) itself (Mattia & Del Giudice, 2002). This entails that our MUA is proportional to *ν*(t), i.e. to the firing rate of the sampled cortical volume. The aggregate spiking activity of neuronal populations in the vicinity of the electrode, without spike detection, was estimated with similar methods in previous work (Choi, Koenig, Jia, & Thakor, 2010; Mattia et al., 2013; Sharma et al., 2015; Stark & Abeles, 2007). As a last step, logarithmically scaled MUAs were smoothed by a moving average (40 ms window, 5ms step which led to a time series of continuous values with a dt=1 ms).

### Quantifying information dynamics with Transfer Entropy

Based on the MUA obtained from each electrode site, we used multivariate **Transfer Entropy** (*TE*) (Komárek, Hrnčíř, & Štěrbová, 2001; Schreiber, 2000) to infer patterns of local information dynamics in the PMd, the directed functional relationship between different neural populations (modules) and the collective state of the information flow through the network at the time of the motor decision. Recent and thriving literature (Bossomaier, Barnett, Harré, & Lizier, 2016; Goetze & Lai, 2019; Ito et al., 2011; Michael Wibral, 2014; Newman, Varley, Parakkattu, Sherrill, & Beggs, 2022; Ramos, Macau, Lopes, & Machado, 2017; Vicente, Wibral, Lindner, & Pipa, 2011; Wibral et al., 2011; Wollstadt, Martínez-Zarzuela, Vicente, Díaz-Pernas, & Wibral, 2014; Xiong, Faes, & Ivanov, 2017) has shown that *TE*, particularly in its multivariate formulation, is more accurate than other measures (Novelli & Lizier, 2021; Ursino, Ricci, & Magosso, 2020), also capturing the effect of nonlinear interactions in the system dynamics without assuming any generative model. Given an ensemble of M time series, the multivariate information transfer from a time series *X* to a series *Y*, conditioned to the remaining *Z*_*k*=1,..,*M*−2_ series, can be quantified considering the present values of *Y* and the past values of both *X* and the *Z* through (Faes, Kugiumtzis, Nollo, Jurysta, & Marinazzo, 2015; Montalto, Faes, & Marinazzo, 2014; Xiong et al., 2017):

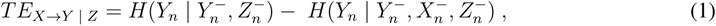

where *Y_n_* is the vector that represent the present state *n* of 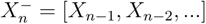, 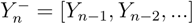 and 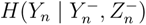 are the vectors that represent the past values of *X*, *Y* and *Z* respectively. The vertical bar stands for conditional probability, e.g. 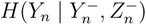 is the entropy of the present state of Y conditioned to the knowledge of the past of *Y* and to the past of the remaining *Z*. *H* is the Shannon entropy (Shannon, 1948), which in the case of *Y* is given by:

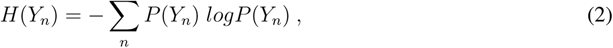

where P indicates the probability density associated with the vector *Y_n_*. Using equation 2 expression 1 becomes:

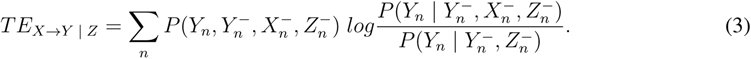

*TE* is *≥* 0 and increases if the history of *X* gives more information about the present of *Y* than the past of *Y* or any other series contained in *Z*. Because the past of *X* is used to predict the present of *Y*, *TE* is not symmetric (i.e. *TE_X→Y | Z_≠ TE_Y →X | Z_*) and defines an information transfer direction, which corresponds to a driver-to-target causal relationship. Approximating the vectors that represent the time series’ history, a process known as *embedding*, is critical when estimating *TE*. The optimal embedding should include the 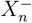, 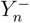 and 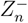 components that are most informative in describing *Y_n_* (Montalto et al., 2014). We used a non-uniform embedding scheme (Faes, Marinazzo, Montalto, & Nollo, 2014) and computed *H* with kernel estimators (for a detailed description of embedding methods and estimators for computing *H*, please refer to the works of Faes and colleagues and the references cited therein (Faes et al., 2015, 2014; Faes, Nollo, & Porta, 2011; Xiong et al., 2017)). Non-uniform embedding is a more reliable alternative to uniform embedding, which would choose the previous values 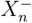, 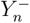 and 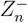 *a priori* and separately for each time series (Montalto et al., 2014; Xiong et al., 2017). Instead, non-uniform embedding selects past components based on maximum relevance and minimum redundancy requirements, gradually removing non-informative terms. Maximum relevance means the most significant predicted information components. If the non-uniform embedding procedure selects at least one component from the past, the resulting *TE_X→Y | Z_* is *>* 0 and statistically significant. Otherwise, if no component from the past is selected, *TE_X→Y | Z_* is deemed to be non-significant and equal to 0 (Montalto et al., 2014). In the selection of the past values, three parameters are important. The maximum number of prior values, or maximum lag *l*; the embedding lag *τ*, that gives the number of steps to take into the past and the offset *u* from where to start to count the steps (e.g. from the first past value, second etc..). In non-uniform embedding methods, the parameters are initialized at the start of the trimming procedure and then sequentially tested for significance. To avoid any bias due to an arbitrary choice of the past values, we initially fixed a “relaxed” uniform embedding vector containing the past up to *l*=50 ms with *tau* and *u* set to 1. Then, we let the non-uniform embedding procedure prune the non informative terms and find the optimal embedding. The statistical significance of past values was progressively assessed by cycling through the past components (up to *l*) and comparing them with a null distribution built from empirical data using a time shifting randomization (Faes, Nollo, & Chon, 2008; Faes et al., 2011; Montalto et al., 2014; Quian Quiroga, Kraskov, Kreuz, & Grassberger, 2002; Varley, Sporns, Schaffelhofer, Scherberger, & Dann, 2023). The components of the past of *X*, *Y*, and *Z* were retained if significant at *α <* 0.01 after 300 shuffles. As expected, the procedure selected only a recent past (no older than 10ms for each *n* of equation 2), aligning with findings from similar studies (Antonello et al., 2022; Shimono & Beggs, 2015; N. M. Timme et al., 2016; Varley et al., 2023; Wollstadt et al., 2014). This is consistent with prior research, which indicates that neuronal populations in the cortex typically establish causal interactions within a time range of 1 to 20 ms (Barthó et al., 2004; Mason, Nicoli, & Stratford, 1991; Swadlow, 1994). To estimate the probability densities necessary for the computation of *H*, we employed the Heaviside Kernel estimator. This estimator reconstructs the P at each data point by assigning weights based on the distance *r* from the reference point *x_n_* to any other point in the time series. Such local exploration of the state space within the neighborhood points of the time series, makes it more robust to inaccurate estimations compared to other approaches (Xiong et al., 2017). *r* is usually chosen to be a fraction of the time series variance (Gómez et al., 2014; Richman & Moorman, 2000; Xiong et al., 2017) so that the dependencies of the estimated entropies from the amplitude of the process are removed. To measure the distance we used the Chebyshev distance (or maximum norm) which is the most commonly used metric (Xiong et al., 2017). The Heaviside Kernel with the Chebyshev distance is equivalent to the sample entropy (Richman & Moorman, 2000; Xiong et al., 2017) which is an established measure to infer complexity of dynamical systems from time series data (Grassberger & Procaccia, 1983; Pincus, 1991) that has been recognized to reduce the estimation bias of conditional entropies. We built single-trial information transfer networks for each behavioral condition of each recording session by customizing some functions of the Matlab MuTE toolbox (Montalto et al., 2014).

Since the application of information theory techniques in neuroscience is still in its early stages, it is worth briefly examining their use on invasive neurophysiological data. Gomez-Herrero (Gómez-Herrero et al., 2015), Bossomaier (Bossomaier et al., 2016), Wibral (Wibral et al., 2011), Newman (Newman et al., 2022) and colleagues have published technical discussions on this topic. Timme and Lapish landmark review (N. M. Timme & Lapish, 2018) is recommended for a full treatment of the critical issues in the field. As summarized in the latter, the main assumption made by experimenters concerns the stationarity of the neural processes under examination. Stationarity of a process means that the probability density function that generates the observable variables does not vary over time, and each observation contributes to the same probability distribution. In practice, this translates into viewing the experimental trials as multiple realizations of the same underlying mechanism. In other words, assuming stationarity implies that each observed realization helps to define the same underlying probability distribution, which finally becomes stable over time, i.e., the lifetime of the experiment (Wollstadt et al., 2014). In a behavioral neurophysiology experiment, the observations correspond to neural activity values, the realizations to repeated trials, and the probability distribution to the probability distribution characteristic of the investigated neural process that is supposed to generate the observed behavior. It is critical to remember that in any experimental paradigm, this distribution is not known a *a priori* and is almost never accessible, in contrast to any theoretical study in which time series are generated based on many repetitions of processes whose analytical details are known. Furthermore, stationarity does not imply a flat variability across realizations, and some variance is expected, reflecting the natural fluctuation of the underlying (unknown) probability distribution (Wollstadt et al., 2014). For these reasons, when an experiment has a multi-trial structure, stationarity over realizations is assumed by default, provided that no slow drifts or trends affect the signal, being the latter one of the non-stationarity factors that most impinge on a correct estimation of *TE* (Xiong et al., 2017). These could arise during training or in an experiment that encourages learning, where the probability distribution of both the brain signal and the behavioral parameters changes over time by definition (Wollstadt et al., 2014).

As a first step in using *TE* on our data, we analyzed the single-trials MUA profiles of each module. Our estimate of MUA is baseline corrected so any potentially distorting slow trends are eliminated. In addition, to remove noisy outliers from our data, we excluded from the analysis the trials for which the MUAs showed peaks with an amplitude that exceeded the average of the activity by 2 standard deviations in any of the time bins of the epoch of interest and for over 80% of the channels (Bardella et al., 2020) (which accounted for less than 2% of the total number of trials). This ensures that artifacts caused by non-physiological peaks and/or segments of ulta-rapid oscillations of high variance are excluded from the analysis. These non-stationary artifacts are among those that have the most impact on trustworthy *TE* estimation since they could influence the probability distributions in equation 3 in such a way that outliers would dominate most of the signal variance, significantly biasing the estimation of the temporal link between the current and past states of the time series (Xiong et al., 2017).

### Graph theoretical analysis

We obtained a single-trial *TE* matrix in the epoch T for each behavioral condition (Go, correct Stop trials and wrong Stop trials) for both monkeys. Assuming that the nodes are single modules (discrete neuronal populations producing MUAs) and the weighted edges are the *TE* values, we interpreted the *TE* matrix as the adjacency matrix of a directed weighted network. We disregarded the main diagonal of the matrix thus excluding self-loops from the networks. Then, we aggregated the trials of the same condition for each recording session to get the appropriate trial-averaged matrices on which we applied **percolation** (Bardella, Bifone, Gabrielli, Gozzi, & Squartini, 2016; Bardella et al., 2020; Bordier, Nicolini, & Bifone, 2017; Bordier, Nicolini, Forcellini, & Bifone, 2018; Callaway, Newman, Strogatz, & Watts, 2000; Del Ferraro et al., 2018; Gallos, Makse, & Sigman, 2012; Mastrandrea et al., 2017, 2021; Nicolini & Bifone, 2016; Nicolini, Bordier, & Bifone, 2017; Vlasov & Bifone, 2017). During percolation, the iterative pruning of links based on their weights is used to monitor the stability of network topology. Percolation was used in this work for two reasons. First, to remove weaker links that were influenced by the statistical noise of trial averaging. Indeed, false positives *TE* values may emerge from the constraints of limited sample size, even when no direct link exists between the variables (Bossomaier et al., 2016; Novelli & Lizier, 2021). Our pipeline included percolation as a topology-based data-driven method for an additional fine-grained cleaning. This also remove the hindrance of optimal thresholding, a common problem when dealing with connectivity matrices (Bardella et al., 2016, 2020; Nicolini & Bifone, 2016), without the need for computationally intractable full error-correction inference of each link at the whole network level. Second, percolation was used to analyze how hierarchical information flow between nodes shapes the complexity of network topology. Percolation analysis produces a curve that plots the size of the largest **connected component** of the network as a function of link weights. Since the networks we studied are directed, the definition of connectedness for directed graphs must be considered. Without delving into much detail, two definitions for a giant component of a directed graph can be given: giant strongly connected component (GSCC) and giant weakly connected component (GWCC) (Graf, 2016; Squires et al., 2013). Very briefly, a GSCC is a subgraph in which every vertex is reachable from every other vertex following the direction of the links. Instead, in a GWCC this is not possible. In other words, in a GWCC a path between every two vertices exists only in the underlying undirected graph. The GSCC thus reveals the network configurations in which information is exchanged cyclically through paths connecting all nodes. We have previously shown that this configuration is important especially during Stop trials, when the PMd network is expected to express quantitative contributions to the inhibition process (Bardella et al., 2020). Hence, we used the GSCC. For simplicity, in the Results we will refer to the threshold at which the fragmentation of the GSCC begins (formally, the percolation threshold) as the ”descent point” of the curves. Such a threshold achieves the best balance between the removal of spurious links and the consequent loss of information potentially encoded in those links (Bordier et al., 2017; Nicolini et al., 2017). The power of percolation lies in displaying the stepped pattern of the curve, which suggests a transitory stable network configuration composed of hierarchical connections between nodes. A steeper curve with no or few leaps, on the other hand, indicates a single incipient component that dominates the network and fragments swiftly in fewer or a single step. This process naturally discloses simultaneously both network robustness and its hierarchical self-organization. Notably, the percolation threshold identifies the level of complexity of the network: a higher threshold means a stronger component; a stronger component means that more links need to be removed to fragment it; more links implies more paths between nodes and thus higher complexity. Thus, we could estimate the complexity of information flow-paths within the network (under different behavioral conditions) by evaluating fragmentation at higher thresholds (see (Bardella et al., 2016, 2020; Del Ferraro et al., 2018; Gallos et al., 2012) for additional details on the percolation analysis applied to brain networks). On each trial-averaged matrix, we measured the Vertex Degree (VD), which is the number of links to each node *i*:

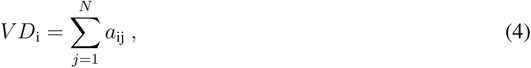

where *a*_ij_ is the generic entry of the adjacency matrix and N is the number of nodes. In directed networks one can distinguish between in-degree (the number on inward links) and out-degree (the number of outward links). For each behavioral condition of each recording session, we estimated the in/out-degree probability for both animals (see supplementary Figure S1). If the variance of the degree distribution is significantly larger than its mean, tails in the distribution arise and network hubs are found. Hubs are thus nodes that significantly exceed the average degree of the network (here we used mean *±* 2 standard deviations of the out-degree distributions (Albert & Barabási, 2002; Barabási, 2013; Barabási & Albert, 1999)). After hubs were identified, we sorted nodes into clusters to better understand their contribution to information transfer. We labeled the nodes based on the two possible outputs of the motor behaviors studied in the task: movement generation (correct Go trials) and movement inhibition (correct Stop trials). To summarize the total quantity of information exchanged between clusters we defined an interaction measure *I*:

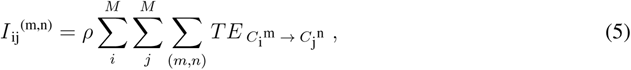

where C is the module’s cluster, M is the number of clusters and m and n run over the all possible combinations of nodes within each cluster. 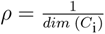 is a normalization factor that accounts for the heterogeneous number of nodes within each of the clusters. Thus, our *I*_ij_ is equivalent to a node strength computed on a renormalized graph formed by M clusters. The strength of a node in a weighted graph is the total of the weights of the links that connect to the node. The values and directions of *I* reflect the cluster’s location in the network communication hierarchy. We enclosed all of the interactions defined by the empirical *TE* matrix in a MxM matrix, with each node representing a functional cluster. *I* was calculated for each recording session and behavioral condition, and the sessions-averaged element (mean *±* SEM) 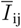 was derived (see Figure 7 and Tables S2 and S3).

### A null model

We built a null network model to assess the statistical significance of the results beyond what was attained by the randomization approach used in the *TE* computations. As noted in a recent work (Cimini et al., 2019), selecting a suitable null model remains a thorny issue in network science. In this study, we drew our conclusions exclusively in a data-driven fashion, with no other constraints. Thus, the simplest null hypothesis is that the results are caused by non-trivial temporal dependencies among neural activities and cannot be explained by the distribution of MUA values and the length of the time series alone. To test this, given a recording session and a behavioral condition and trial, we generated a synthetic time series for each recording site (node of our *TE* network) by randomly permuting the corresponding MUAs. In this way each surrogate time series has the same basic statistical properties of the original one (mean, variance and histogram distribution), while relationships across the series of different nodes are destroyed (Faes, Pinna, Porta, Maestri, & Nollo, 2004). For each surrogate single-trial graph we then preserved the link density of the corresponding empirical matrix and computed the trial-averaged null *TE* network. Repeating the procedure 500 times for each condition and recording session, we obtained an ensemble-averaged *TE* network with which we compared our empirical graph metrics (see Figure S2). We chose this null model because it is the least restrictive and no assumptions are made about the connection patterns and weight distributions of the synthetic networks. This was in line with our goal: to test that the information patterns in the network could not be detected solely by constraining the activity distribution at the node level.

## RESULTS

Table S1 shows behavioral data from the 21 recording sessions used in the study, 12 from Monkey P and 9 from Monkey C. In order to compute the SSRT, we ensured that the race model’s fundamental premise was followed in all sessions. To be more specific, the Go process in the Stop trials must be the same as the Go process in the Go trials to compute the SSRT. This assumption is usually satisfied by showing that RTs in wrong Stop trials are shorter than RTs in Go trials, as shown in Table S1. This indicates that all trials are drawn from the same distribution of RTs. Across all sessions, the average RT for Monkey P was 656 ms (SD = 129 ms) and 632 ms (SD = 119 ms) for Monkey C. The average SSRT was 217 ms (SD = 34 ms) for Monkey P and 213 ms (SD = 26 ms) for Monkey C.

### Neural activity significantly contributes to motor control at the single-site level

Fig 2 illustrates the time course of the average MUA obtained from each cortical site separately during Go trials (dark green traces), correct Stop trials (red traces), and wrong Stop trials (dark blue traces) during the time (epoch T in Figure 1) preceding the movement onset (in Go trials and wrong Stop trials) or the end of the SSRT (in correct Stop trials) for one example session of Monkey P (S6 of Table S1). The activity in the wrong Stop trials appears to be indistinguishable from the activity in the Go trials for all sites. In contrast, the presentation of the Stop signal (red vertical line) and the effective cancellation of movement produce a distinct pattern in many of the locations, with the spiking activity modulated (increasing or decreasing) compared to the activity during movement generation. In this specific session we found a significant difference between the Go and correct Stop conditions in 78% of the recording sites. Conversely, there was no significant difference in spiking activity between the Go and wrong Stop trials at any of the sites. A similar trend was found in all recording sessions (Table S4), confirming previous findings on the role of PMd in inhibitory motor control and the relevance of the epoch of analysis in motor decisions during Stop trials (Bardella et al., 2020; Battaglia-Mayer et al., 2014; Giarrocco et al., 2021; Mattia, Ferraina, & Del Giudice, 2010; Mirabella et al., 2011; Pani et al., 2014, 2022, 2018; Ramawat et al., 2022). This result, however, does not provide information on how the nodes in the network interacted in the different behavioral conditions.

**Figure 2.**
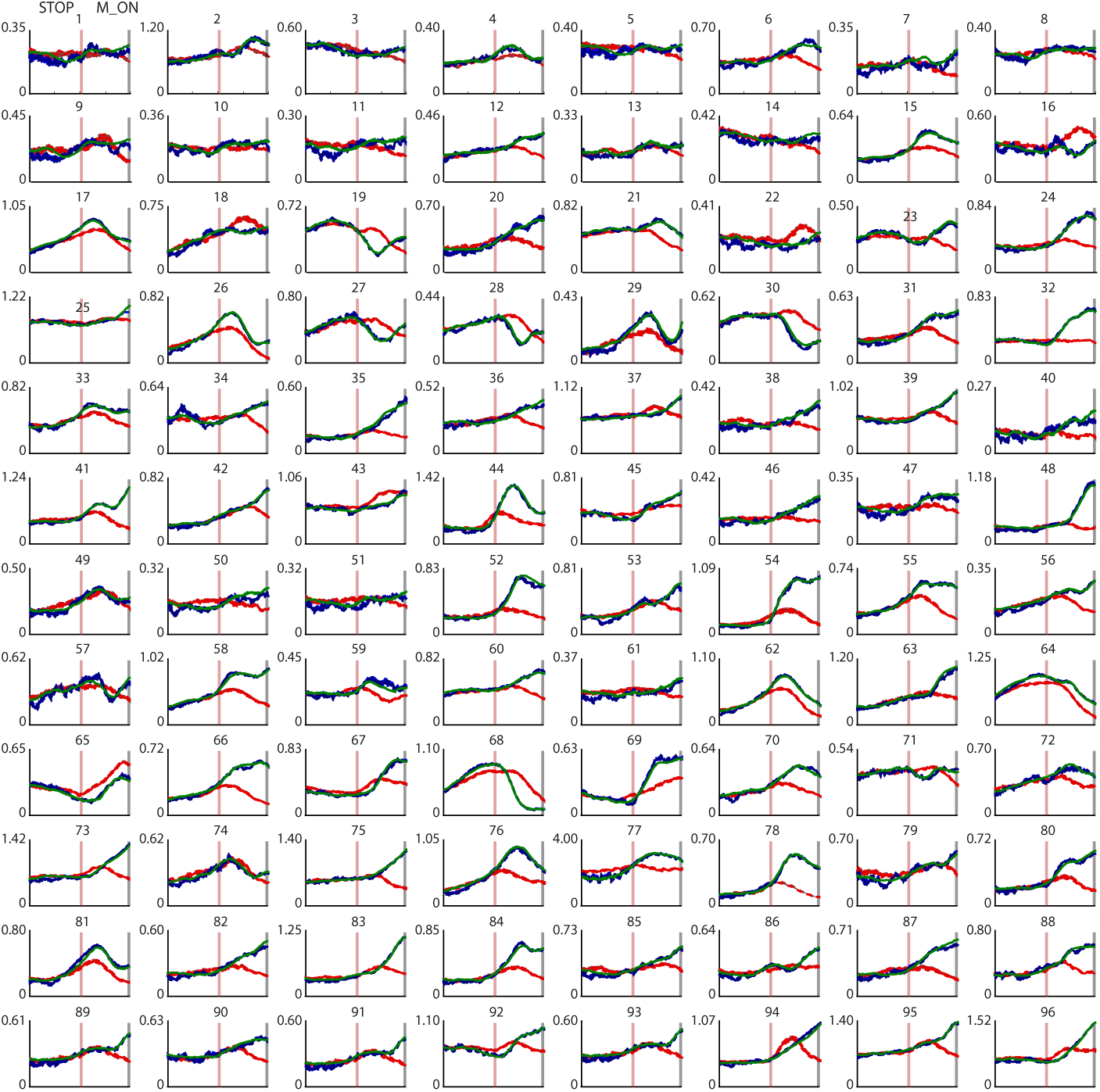
Neuronal activity modulation for different trial types. Neuronal MUAs (mean *±* SEM) in the epoch T for the three behavioral conditions for all recording sites (modules) of S6 of Monkey P. Dark green traces show the average activity during Go trials aligned to the movement onset (gray vertical line). Dark blue traces show the average activity during wrong Stop trials aligned to the movement onset (gray vertical line). Red traces show the average activity during correct Stop trials aligned to the Stop signal presentation (light red vertical line) and plotted until the end of the SSRT. For each plot the Y axes are units of MUA while the X axes mark the time during the epoch T; ticks, every 100 ms, are indicated for panels in the top row only. Numbers above each panel indicate the recording site. The actual spatial mapping of the recording sites is not preserved and are here shown in cardinal order from 1 to 96.

Fig.3 shows the *TE* matrix for each trial type for the data provided in Figure 2. The dense horizontal stripes that change between Go, correct, and wrong Stop trials suggest that, depending on the behavioral situation, a few of the sites might influence the activity of many others during the epoch where the motor choice is taken. In this picture, any recording location is interpreted as a network node, and any significant *TE* value as a link. To describe network topology we first calculated the vertex degree (VD) for the average networks, thresholded via percolation analysis after removing the noisiest links (Bardella et al., 2016, 2020; Bordier et al., 2017; Nicolini & Bifone, 2016; Nicolini et al., 2017) (see Section). A high VD indicates that some modules are functionally related to many others and serve as hubs. Hubs only appeared in the out-degree (*V D_out_*) distributions (see Figure S1 and S2). In each session, we labeled the nodes based on the behavioral condition in which they were hubs. Were classified as **C_1** or **C_2**, the nodes that were hubs only in trials where the movement was generated or successfully canceled, respectively. The **C_3**, included all the nodes that were hubs in both Go and correct Stop trials. The **C_4** included the nodes that were never hubs. Interestingly, no new hubs were found when movements were incorrectly inhibited after a Stop signal (wrong Stop trials). This implies that the same neuronal clusters (groups of nodes) recruited in Go and correct Stop trials are also involved when errors occur in Stop trials. The percentage of nodes belonging to each cluster throughout all sessions is shown in Table S5.

**Figure 3.**
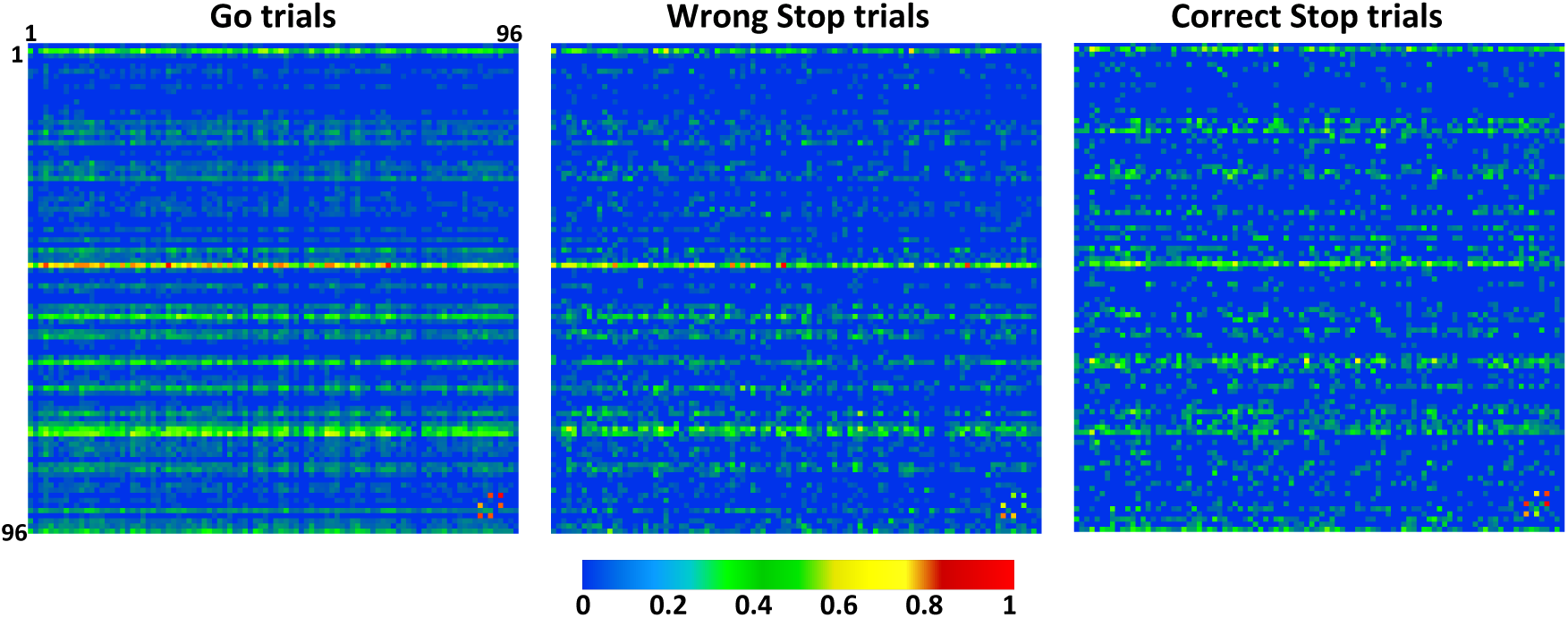
Average. *T E* **matrices** of Go, wrong and correct Stop trials for data in Fig.2. Each entry of the matrix is color-coded according to the corresponding *T E* value averaged over trials. Statistically significant driver *→* target relationships correspond to *T E* values *>* 0 for the pair of modules considered. Here, for graphical purposes only, *T E* is normalized to the maximum across behavioral conditions so to obtain *T E ∈* [0, 1] (color bar). Axes labels indicate recording sites.

One might wonder if there is any internal hierarchy between the clusters, specifically whether the *V D_out_* of the nodes in a cluster changes depending on the behavioral conditions. To answer, we kept the cluster label fixed and studied how the *V D_out_* of the nodes in each cluster differed across trial types. **C_3** consistently exhibited the highest *V D_out_* values across all three behavioral conditions across all sessions (see Table S6), indicating that they are the primary hubs for the spreading of information (Kruskal-Wallis, *p <* 0.01, Bonferroni corrected for multiple comparisons). Thus, **C_3** are at the highest hierarchical level in the network because they regulate information transfer regardless of whether the decision is to move or stop. On the other hand, **C_1** and **C_2** have distinct levels of engagement, with lower *V D_out_* values in Go and correct Stop trials, respectively, shifting their role based on the type of motor choice being processed. In Go trials, the **C_2** had lower *V D_out_* values than the **C_1**, whereas in correct Stop trials, the **C_1** had lower *V D_out_* values than the **C_2**. Interestingly, the **C_1** had larger *V D_out_* values during wrong Stop trials than during correct Stop trials, whereas the **C_2** had the opposite (Kruskal-Wallis, *p <* 0.01 Bonferroni corrected for multiple comparisons). This reveals that during wrong Stop trials, the neuronal processing of information is midway between Go and correct Stop trials, but more comparable to Go trials. The presence of four distinct clusters with varying *V D_out_* shows that some nodes serve as driving hubs only in relation to a specific behavioral outcome. Such heterogeneity implies that the patterns of information transmission are fine-grained and are adjusted according to whether the planned movement is executed or withheld. The topology of the PMd network for data in Fig.3 is depicted in Fig.4, highlighting the **C_1** and **C_2** hubs, that change between conditions (for the networks including also the **C_3** see Figure S4; see Figure S5 for examples from other sessions). The information network has a centralized organization in Go trials (and to a lesser extent in wrong Stop trials), corroborating our previous study that showed how such functional conditions facilitate movement generation Bardella et al. (2020). Fig.4 (and Supplementary Figures S3 and S4) show how the **C_1** act as hubs during Go and wrong Stop trials, whereas the **C_2** are hubs during correct Stop trials only. The optimal centralized transmission present during Go trials is disrupted during correct Stop trials, where the links arrangement is more complex and compounded, with more intersections and less directness compared to Go trials. This suggests that the network retains its core feature (the existence of hubs) across trials but varies in the way they are overall deployed based on the required behavioral control.

**Figure 4.**
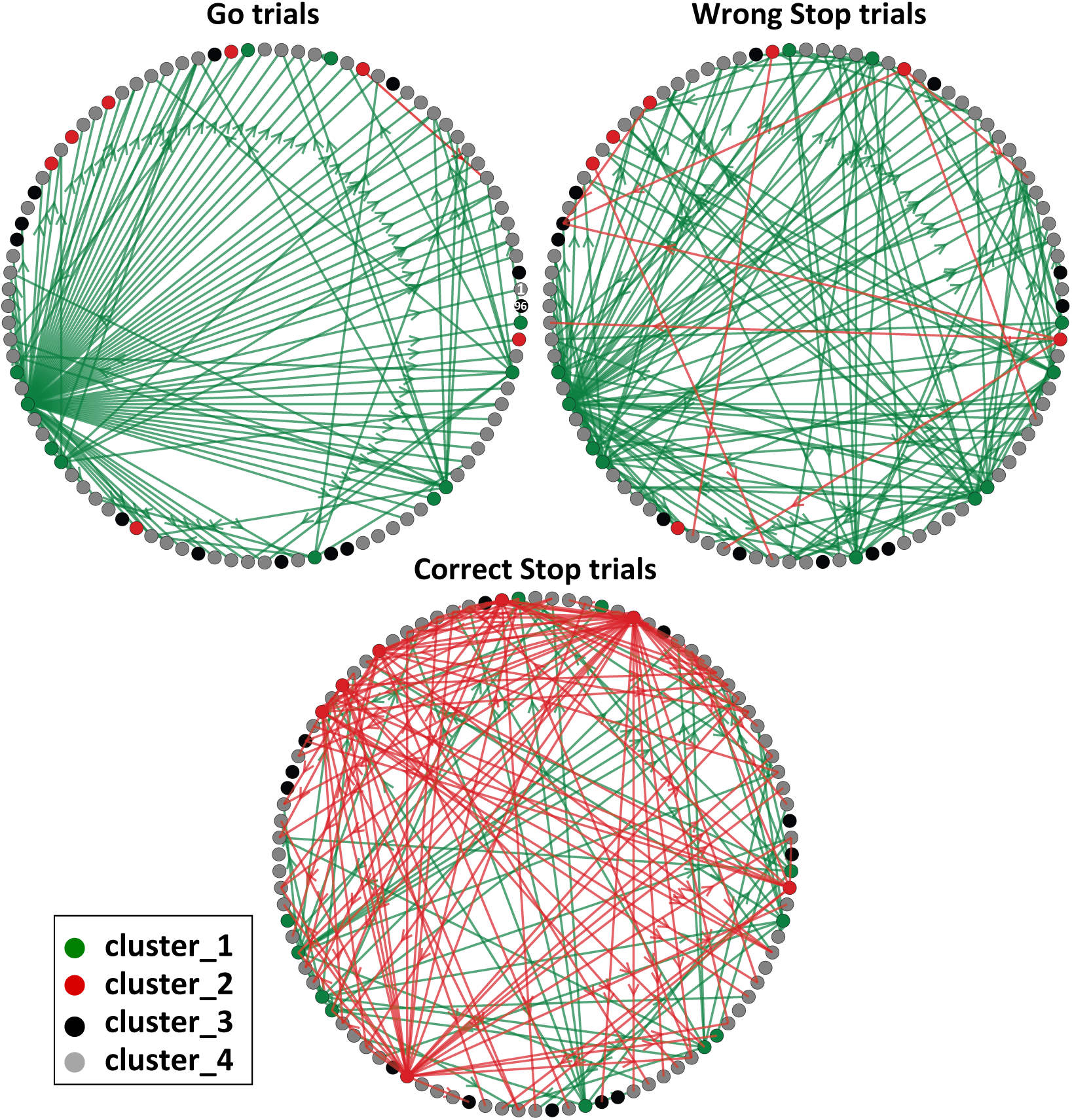
Information network of Go, wrong, and correct Stop trials. Data from the same recording session of Fig.2 and Fig.3. Each link is a significant *T E* value (i.e., a non-zero entry in the *T E* matrix). The number of outgoing connections of each module reflects its *V D_out_*: the number of modules on which it acts as a driver. Each link is color-coded according to the cluster to which its source node belongs. Information-spreading hubs are observable. Here, for illustrative purposes and to better highlight the differences between behavioral conditions, the links of the C_3 nodes are removed and for each network only the 20% of the strongest links are shown (for the complete version including the C_3 see Figure S4). Recording sites 1 and 96 are marked in the Go trials graph. The coordinates of each node on the circle are preserved across behavioral conditions.

### The PMd network becomes more complex in Stop trials

To better understand how the hierarchical relationships between the clusters relates to the overall network organization, we analyzed the *V D_out_* as a function of *TE*. This involved tracking the changes in *V D_out_* of each cluster as the percolation process progressed. In this way, the resilience of the information carried by each cluster is compared to the information spread by the others. We expect that a cluster that is functionally relevant to a specific behavioral condition should possess not only a large number of connections (a high *V D_out_* value) but also a substantial number of these connections should exhibit high and robust information content (reflected in high *TE* values). Results are shown in Fig. 5 as the average of all recorded sessions of each animal. The analysis confirmed the **C_3** as the highest hierarchical level cluster in all three behavioral conditions. The highest values of *V D_out_* as a function of the amount of information processed, indicate that it had the most links with high and robust information content. The **C_1** and **C_2**, on the other hand, were placed between the **C_3** and the **C_4**, and their hierarchical level changed depending on the trial type. Thus, the network’s global structure is similar in both wrong and correct Stop trials. Inspecting the percolation curves we could infer the level of network complexity under the three behavioral settings (see Section). Figure 6 shows that the percolation curves of the Go trials evolve differently from those of the Stop trials (whether correct or wrong) and the null model for all sessions. The curves for the Go trials begin their descent at lower TE values and have a more stepped pattern (normalized among conditions). Therefore, the Go trials have fewer paths belonging to the giant component itself, with most links with high information content heading outwards to the rest of the network. This is a symptom of an ideal network setup with more direct and organized communication than in the Stop trials. One important implication is that in the Go trials the number of redundant links is very low. In other words, there are just a few links that deviate from the main common structure. During Go trials most high-weight links are used for direct communication between hubs and the rest of the network, lowering the level of complexity compared to Stop trials. Conversely, in Stop trials, a complicated rearrangement happens, with denser communications and high-content information traveling more within the large component than outwards. The proximity of the Stop trial curves to the null model (NM) reveals that, although they differ from a purely random situation, they still adhere to some of its communication rules. Indeed, in the NM, which is inherently complex, one would anticipate a configuration where links are randomly arranged regardless of their weights. This results in a network that is dense with paths between nodes but lacks stable configurations, as indicated by the gray curve’s abrupt descent without intermediate steps. Most notably, such a network would lack hubs (see also Figure S1). The slope of the curves in panel c demonstrates that the quantity of information necessary to obtain equivalent stable configurations (i.e.the same values of y in panel a) is smaller in Go trials than in Stop trials. This suggests that the higher inherent complexity of the latter is related to increased resource recruitment. These findings confirm how the local links of the clusters impact the overall state of the network, changing how it manages information transmission.

**Figure 5.**
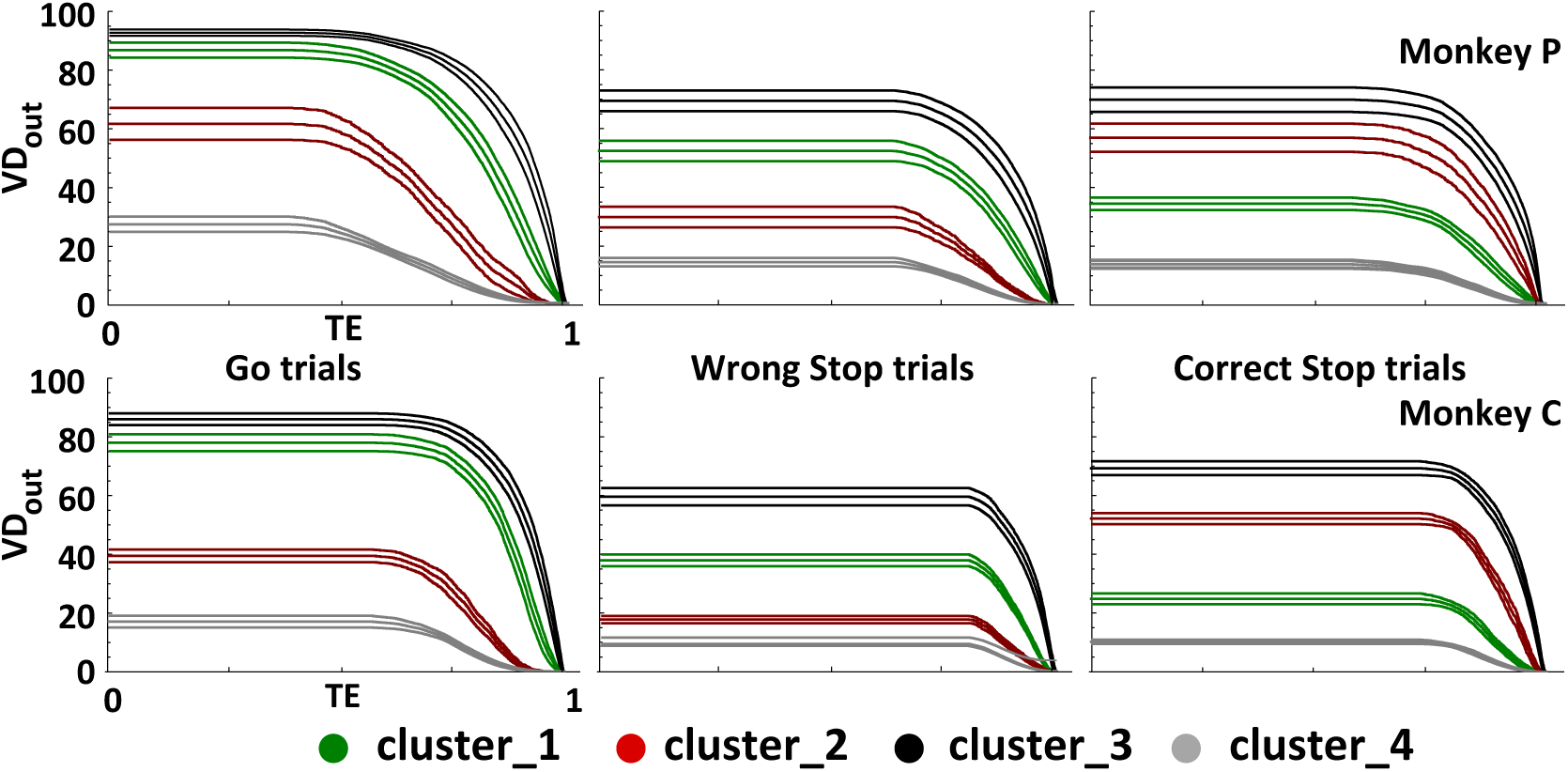
Clusters robustness analysis. Robustness is inferred by monitoring the evolution of *V D_out_* (y axis) as a function of the information processed (*T E*, x axis). Panels show the robustness curves averaged (mean *±* SEM) over recording sessions compared between clusters and across behavioral conditions. Curves are color coded accordingly to the corresponding cluster as in Fig.4.

**Figure 6.**
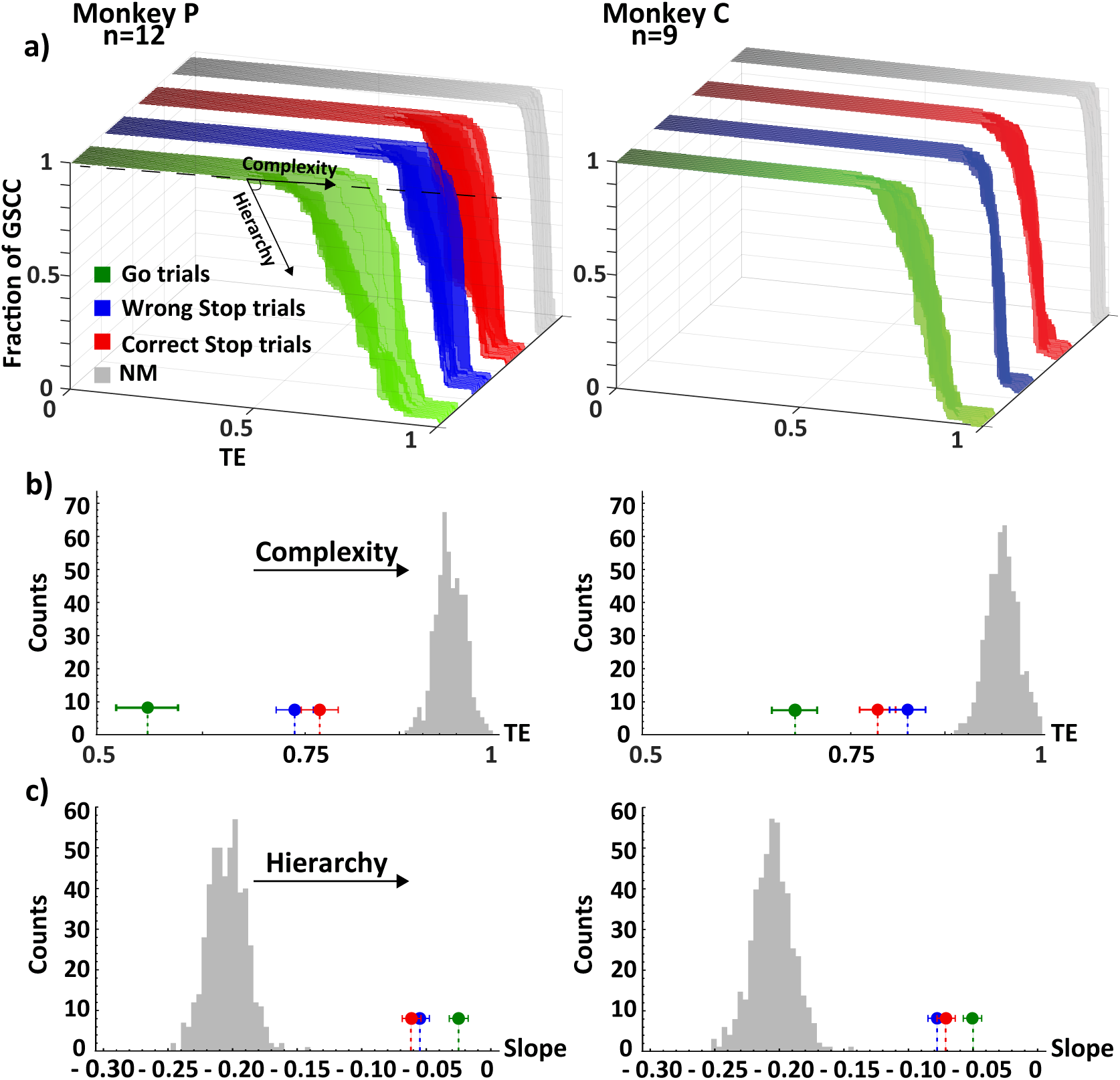
Percolation analysis. **a:** percolation curves over recording sessions compared across behavioral conditions and vs the null model for both animals. Each curve is a recording session. As sketched, the level of complexity of the underlying networks increases traveling the curves from left to right, whereas the hierarchy can be explored climbing down the steps of the curves (the fraction of GSCC, z axis), measured in units of information (*T E*, x axis). **b:** comparison between the point of descent between behavioral conditions (mean *±* SEM) and against the null model distribution. **c:** comparison between the measured slopes between behavioral conditions (mean *±* SEM) and against the null model distribution. Stop trials resulted more complex and less hierarchical than Go trials while having comparable values between themselves (Kruskal-Wallis, *p <* 0.01 Bonferroni corrected for multiple comparisons). In all panels, for simplicity and homogeneity, TE is reported normalized to its maximum so to obtain TE *∈* [0, 1]. The null distribution is obtained from 500 runs of the null model (see Section). Since the null distributions for the different behavioral conditions were indistinguishable, we only reported one for brevity. (**GSCC**: Giant Strongly Connected Component; **NM**: Null Model).

### A recap

In summary, we have demonstrated the existence of diverse neural modules/clusters (Figure 4) and have described how these clusters interact and distribute information in various ways, thus contributing to the state of the mesoscopic network and its level of complexity, each with its own distinct features (Figures 5 and 6). The wiring of information transmission within the network can be efficiently summarized in Figure 7 using the interaction measure *I* (see Materials and Methods, Table S2 and S3 for further details). The **C_3** cluster is the primary determinant of the global network state, while the **C_1** and **C_2** play supporting roles. The strength and direction of the interactions between **C_1** and **C_2** hint at the specialization of these clusters for behavioral conditions. Specifically, in Go trials, information travels from **C_1** to **C_2** while in correct Stop trials from **C_2** to **C_1**. The flow of information from the **C_1** and **C_2** to the **C_3** followed the same pattern. The network topology during wrong Stop trials was, as expected, a hybrid of the configurations observed in Go and correct Stop trials. We discovered that correct and wrong Stop trials have the same overall topological organization but differ in the details and hierarchies of information processing. In Wrong Stop trials, we observed a mixture of interactions between clusters, similar to what was observed in Go trials, along with evidence of attempted inhibition. Our findings revealed that motor control at the PMd level revolves around a well-defined hierarchical scheme in which **C_3** are the pivotal nodes in the network and the **C_1** and **C_2** provide critical support in movement generation and inhibition, respectively.

## DISCUSSION

In this work, we investigated the patterns of local information transfer in a cortical network directly involved in movement decision-making (the PMd). We used a combined *TE* and graph theory-based approach to analyze the aggregate spiking activity of simultaneously recorded populations of neurons. Our results contribute to moving forward the knowledge on the neural basis of motor control at different levels.

### A topological approach to local information propagation in a motor decision task

Although *TE* is growing in popularity in modern neuroscience, its application with graph theory to study patterns of directed connectivity has been limited, and restricted mostly, to either single neurons (Gerhard, Pipa, Lima, Neuenschwander, & Gerstner, 2011; Shimono & Beggs, 2015; Varley et al., 2023) or in vitro (Buehlmann & Deco, 2010; Kajiwara et al., 2021; Newman et al., 2022; Nigam et al., 2016; Orlandi, Stetter, Soriano, Geisel, & Battaglia, 2014; Shimono & Beggs, 2015; Thivierge, 2014), and in silico studies (Ito et al., 2011). Its applications to behavioral neurophysiology have been instead extremely limited. However, in recent times, authors have started to explore more consistently the joint information theory-complex networks approach under various experimental settings (Kajiwara et al., 2021; Newman et al., 2022; Schroeder et al., 2016; Shimono & Beggs, 2015; N. Timme et al., 2014; Varley et al., 2023). In an early contribution, Gerhard et al. (Gerhard et al., 2011) examined spike train recordings from a rhesus monkey’s visual system during a fixation task, employing a different approach to measure directed connectivity. Besserve and colleagues (Besserve, Lowe, Logothetis, Schölkopf, & Panzeri, 2015)are credited with applying *TE* to invasive electrophysiological data from the visual system, although without incorporating graph theory techniques. Another significant contribution comes from Honey and colleagues (Honey, Kötter, Breakspear, & Sporns, 2007), who explored the relationship between the functional and structural properties of a large-scale interregional anatomical network in the macaque cortex using *TE*, albeit at different spatial and temporal scales. Wollstadt and collaborators (Wollstadt et al., 2014) considered larger spatial and temporal scales than those investigated here and introduced the “ensemble method” for estimating TE under nonstationary conditions, which utilizes the multi-trial structure in neuroscience data for more accurate time-resolved *TE* estimation. This represents a difference from our work, where, instead, the exchange of information is inferred in a fixed time window. The choice was motivated by our goal: characterize the state of the PMd network at a specific epoch of the task and compare it across behavioral conditions. To do so, we needed several network realizations (over trials) in order to obtain a statistically robust estimate of its topological properties. In contrast, the ensemble approach would have led to a single representation of the temporal evolution of the network. A temporally precise but unique one. It could be argued that this constitutes a limitation of our analysis. Instead, we believe it is the way to answer a different scientific question. We also believe that the ensemble method is extremely powerful and it could be of interest to use it on our data in future studies. An attempt to study voluntary action control through analysis of directed connectivity was made by Jahfari and colleagues (Jahfari et al., 2011) but on human fMRI data. Hence, to the best of our knowledge, this report is one of two studies that use graph theory to analyze the information transfer network of a specific cortical area at the mesoscale level *in vivo* and during cognitive motor control. The only other work on the topic known to us is the extremely recent paper from Varley et al.(Varley et al., 2023) who, using the same dataset as Dann et al.(Dann, Michaels, Schaffelhofer, & Scherberger, 2016), quantified information transmission between single neurons in the fronto-parietal areas AIP, F5, and M1 during various epochs of a delayed grasping task. The authors compared several components of information processing, including also a multi-variate estimate of Transfer Entropy very similar to ours, and inspected how they changed as the task unfolded (see next paragraph). The level of description obtained in our work is more detailed compared to previous ones on motor control. Indeed, we were able to specify how the decision on whether to move or to stop is implemented in PMd at the meso-scale population level, who are the (key) players that manage information transmission, and how information propagation shape the hierarchies and the complexity of the network depending on the behavioral conditions. Notably, in our framework, neither any a priori assumption nor a specific modeling technique was needed. Our entirely data-driven approach, in addition to complementing the most recent models for motor generation and suppression (Boucher, Palmeri, Logan, & Schall, 2007; Lo, Boucher, Paré, Schall, & Wang, 2009) allows us to address their primary limitation, which involves the need to fine-tune numerous biophysical parameters to achieve an accurate fit with experimental data. It is known that to fully understand the neural mechanisms behind motor control future research should focus on cortico-cortical and cortico-subcortical interactions through massive simultaneous recordings. In this scenario, a topological information-based approach is unquestionably necessary and should be favored over other common approaches that rely on the analysis of covariance between neurons or mesoscopic signals(Chandrasekaran, Peixoto, Newsome, & Shenoy, 2017; Churchland et al., 2012; Clawson et al., 2019; Kaufman et al., 2016; Mattia et al., 2013; Pani et al., 2022). On one hand, these methods enable a precise temporal description of neural variability. In contrast, they lack the ability to describe information dynamics between neurons or provide quantitative insights into network topology and hierarchy. Without this spectrum of details the computational strategy underlying motor control (and neural circuitry computation in general) would remain yet elusive.

### Fine-grained information transfer underlies motor decisions in PMd

We found that neuronal activities could be organized in four clusters that actively participate and interact, with distinct roles, both in movement execution and cancellation. This is a step forward in the conceptualization of the neural processes underlying motor control. Indeed, other studies describe movement generation as the result of accumulation processes toward a threshold, or as a transition of neuronal trajectories across specific subspaces. Some studies describe the accumulation process as instantiated by specific neuronal types (as for gaze-shifting neurons in the frontal eye field and in the superior colliculus for saccades). Others consider the neuronal trajectory reflecting the dynamics of the entire neuronal population without considering specific neuronal types. units (Boucher et al., 2007; Kaufman, Churchland, Ryu, & Shenoy, 2014; Lo et al., 2009; Marcos et al., 2013; Pani et al., 2022; Schall, Palmeri, & Logan, 2017; Verbruggen & Logan, 2008; Wei, Rubin, & Wang, 2015) Instead, we showed that information is hierarchically transferred between actors with a well-identifiable role, e.g. the cluster (C_3) acting as the most important hub for information spreading across behavioral conditions. This reflects the presence of high-order complexity at the population level, even in small cortical regions during behavioral control, regardless of the nature of neurons in each cluster, whether excitatory or inhibitory. Our findings clearly show that hierarchical control is not only exerted across brain regions, but it is also implemented locally by specific neuronal clusters (the C_3 cluster in our case) over the others. This is in line with evidence provided in recent years that the brain is hierarchically organized both globally and locally on a spatial (Bardella et al., 2016, 2020; Felleman & Van Essen, 1991; C. Hilgetag, O’Neill, & Young, 2000; C. C. Hilgetag & Kaiser, 2004; Kaiser, 2010; Kaiser, Görner, & Hilgetag, 2007; Sporns, Honey, & Kötter, 2007; Sporns, Tononi, & Kötter, 2005; Tononi, Sporns, & Edelman, 1994; Varley et al., 2023; Watanabe et al., 2014; Zamora-López, 2010; Zeki & Shipp, 1988) (for a detailed review see Hilgetag et al., 2020(C. C. Hilgetag & Goulas, 2020)) and temporal scale (Cirillo, Fascianelli, Ferrucci, & Genovesio, 2018; Fox et al., 2005; He, 2011; Morcos & Harvey, 2016; Shine et al., 2016; Vidaurre, Smith, & Woolrich, 2017; Zalesky, Fornito, Cocchi, Gollo, & Breakspear, 2014). We revealed both condition-specific and nonspecific neuronal clusters, demonstrating that information transfer and network complexity depend on behavioral outcomes. Our depiction aligns with the current view by identifying specific clusters primarily involved in the generation (C_1 cluster) or inhibition (C_2 cluster) of movements. Additionally, it introduces a high-order level of control (C_3 cluster) not previously proposed in other works.This fully portrays the interaction patterns underlying inhibitory control at the PMd level (see Fig. 7). How does this relate to the current behavioral models? According to the race model, the Stop process should be ’activated’ only in the Stop trials following the presentation of the Stop signal. However, increasing evidence (Drummond, Cressman, & Carlsen, 2017; Dunovan, Lynch, Molesworth, & Verstynen, 2015) suggests that the Stop process is also active in Go trials. Our results show that such co-activation corresponds to information revolving around four clusters arranged in higher-order connectivity patterns. This is testified by the hierarchical levels in which the C_1 cluster and C_2 cluster are ranked in Go and Stop trials. Moreover, our results show that the C_2 cluster exhibits a flow of information directed to the C_3 cluster that is significantly higher during correct Stop trials compared to the other trial types. This suggests that some of the information that the C_2 cluster handles in attempting to abort the movement is directed to influence the C_3 cluster, which are extensively involved in facilitating the overall process. This evidence implies that the reason for the unsuccessful attempts in the wrong Stop trials is associated with the strong flow directed from the C_1 cluster to the C_4 cluster, similar to what occurs in the Go trials This aligns with the limited involvement of the C_2 cluster, which distributes significantly less information compared to properly inhibited movements (see 7 and Tables S2, S3 and S6). The found subdivision in clusters and the presence of hubs reveal that the cortical information dynamics behind motor control is extremely rich and must be accounted for to generate more comprehensive behavioral models. The topological structure detected across behavioral conditions exhibits stable rules for the information transmission: the one based on hubs.This implies that the presence of high-degree nodes is a constituent feature of neural processing in a cortical network directly involved in cognitive control, as is the PMd.. This is consistent with our previous study (Bardella et al., 2020), in which we showed how the functional PMd network organization differs between movement generation and inhibition in terms of hierarchy and centrality of nodes. It is also in agreement with other works that found high degree nodes in silico (Gu, Qi, & Gong, 2019), in cortical and hippocampal in vitro networks (Antonello et al., 2022; Gal et al., 2017; Nigam et al., 2016; Perin, Berger, & Markram, 2011; Schroeter, Charlesworth, Kitzbichler, Paulsen, & Bullmore, 2015; Shimono & Beggs, 2015; N. Timme et al., 2014), in vivo (Dann et al., 2016; Varley et al., 2023) and anatomical structural networks (Honey et al., 2007). Within this topology, different information exchange regimes are realized. On one hand, correct and wrong Stop trials resulted in similar levels of complexity. On the other, during Go and wrong Stop trials clusters were recruited to exchange information in a similar fashion. This indicates a transition of the information network from a centralized state during Go trials to a distinct one during Stop trials. The latter is characterized by higher complexity in information transfer and increased randomness. This change signals a strong association between the network state and the behavioral condition. The structure found in the Go trials implies a high and very robust communication strategy compared to that observed in trials in which a Stop is presented.Conversely, the network configuration associated with a planned and the inhibited movement appears to be topologically very similar to the previous one but with some degree of randomness ’injected’ into the interactions. This shows how a network engaged in a cognitive task can adapt its functionality by altering its complexity and heterogeneity, utilizing different resources while maintaining the stable generating rules of its topology. We interpret the reorganization occurring in correct Stop trials as the execution in the local network, through the intervention of the C_2 cluster, of a command originating from other regions. Indeed, as known, the PMd is part of a larger network subserving motor control based on frontal, parietal, subcortical and spinal cord structures. It is reasonable to think that during Go trials the hubs serve to convey the command to move to other (and possibly anatomically lower) cortical, subcortical, and spinal circuits that will eventually promote muscle activation. What is observed during correct Stop trials could reflect the PMd collective reaction to the incoming inhibitory input that prevents the execution of the programmed movement. The volition to inhibit would be locally implemented as ‘the attenuation of the movement state’,achieved by adjusting the level of randomness in the network’s wiring scheme, resulting in a more complex organization. This appears convenient and straightforward to implement at the network level. To transmit the message, it would only require the recruitment of dedicated resources (the C_2 cluster) and the redirection of information flow through different paths between nodes. The state of wrong Stop trials is topologically related to that of a correct Stop trial and, at the same time, exhibits the same patterns of information transfer as in Go trials. This may result from an erroneous transmission of inhibitory coding. It can be speculated that this is due to a persistent strong recruitment of the C_1 cluster and a contemporary weaker involvement of the C_2 cluster, as it happens during Go trials. This aligns with our recent paper where we analyzed the neural activity dynamics in the same task using a state-space approach (Pani et al., 2022). In that study, we demonstrated the existence of a subspace in which neural activities are confined before movement generation and during movement suppression.

**Figure 7.**
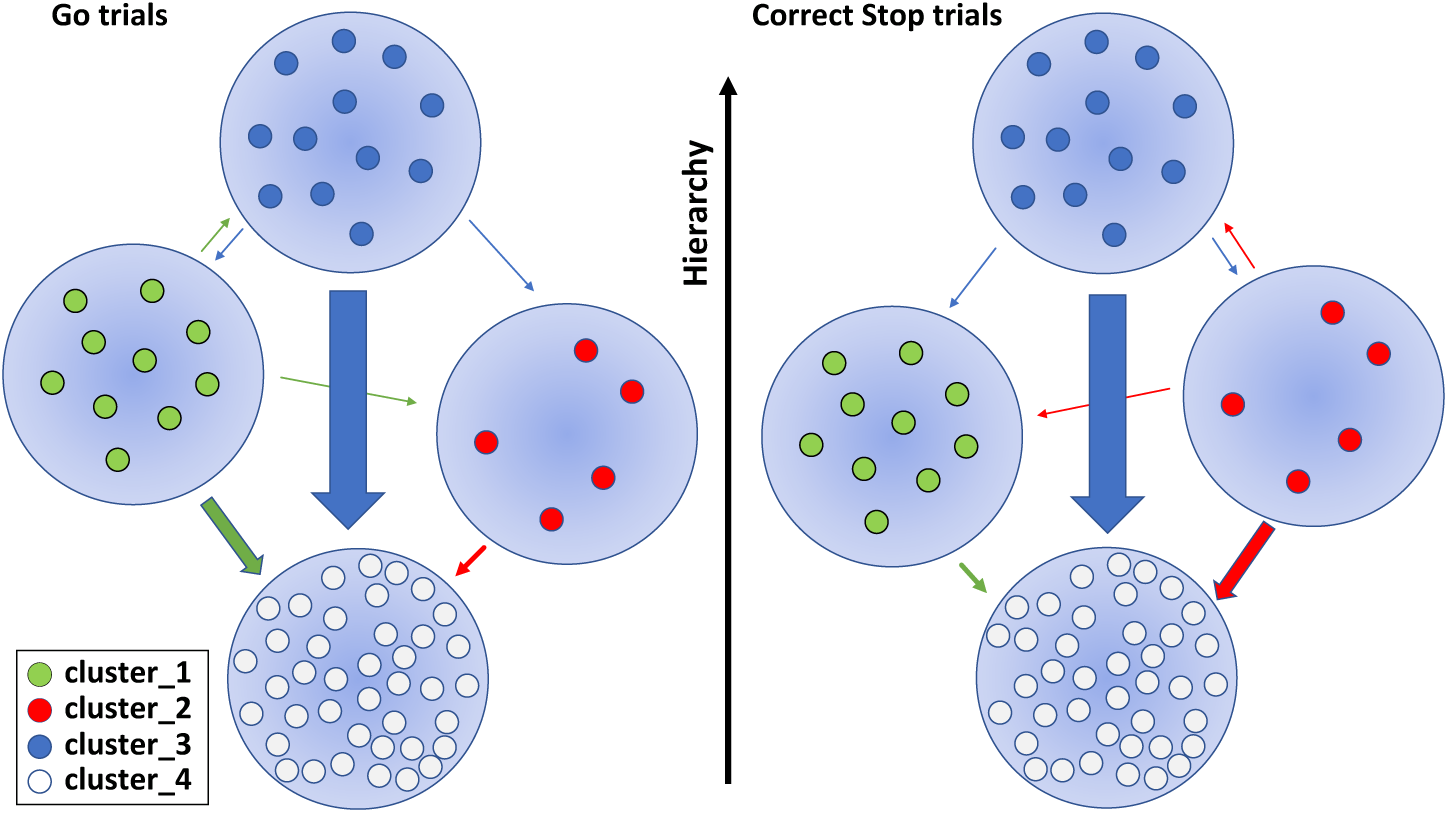
Information spreading among clusters before movement generation and cancellation. Arrows between groups of modules are scaled according to the corresponding average value of **I** (among sessions and animals). The level of hierarchy in which each cluster is ranked is graphically represented as the vertical positioning of the nodes; such level depends on the set of measures explored and could be efficiently summarized by the index I. Since the **C_3** is the dominant cluster in the network, its interactions are always shown. For the other clusters, only the interactions at least one order of magnitude greater than the others and statistically significant across behavioral conditions are shown. Wrong Stop trials are not shown; see Tables S5 and S6 for further details.

One weakness of this study is that we cannot account for the information dynamics between PMd and other structures of the reaching network. Therefore, additional research, in a multi-area framework, will be needed to unambiguously clarify the interactions of the PMd network with other structures and to investigate to whom the clusters project or are connected to. The multi-area approach is instead a strength of the work of Varley’s and colleagues (Varley et al., 2023) with which our results are only partially comparable. They did not record from the PMd and the task they analyzed did not include inhibition trials. For the shared part (the movement trials), the results are in agreement: the presence of hierarchical hubs also emerged as a core topological attribute in their case, further confirming the ubiquitous presence of this feature of neural organization in cognitive control. The ’processing units’ they described, emerge from averaged interactions of macro ensembles of neurons, thus neglecting their decomposition into individual contributions (and their interactions) within the same area. Both works thus have their strengths and weaknesses, testifying how for a more complete understanding of behavioral control this type of approach needs to be systematized. Considering the nature of the mesoscopic signal we studied, it was not possible to resolve the contributions of individual neurons within each of the modules forming the clusters. This relates to another question that remains unexplored in this article: whether and how the excitatory/inhibitory nature of individual neurons could affect cluster composition and role. To this end, the framework of information theory is essential, as extensions of the TE measure have recently been proposed to track the excitatory or inhibitory signatures of information exchange (Goetze & Lai, 2019; Kajiwara et al., 2021). Our conclusions suggest that the collective network organization we discovered represents the neural implementation for the voluntary motor control at the PMd level.

## ACKNOWLEDGMENTS

This work was in part supported by European Union Horizon 2020 Research and Innovation Programme (Grant 945539; Human Brain Project Specific Grant Agreement 3, to SF) and received partial support from the Italian National Recovery and Resilience Plan (NRRP), M4C2, funded by the European Union - NextGenerationEU (Project IR0000011, CUP B51E22000150006, “EBRAINS-Italy”, to SF), from EC ERANET CHIST-ERA 2019 grant ’MUCCA’ (to SF), from grant RM11916B89232364 (to PP) and from grant ’Borse di Studio 2020’ Fondazione Baroni (to GB).

