## Supplementary Material for "Response inhibition in premotor cortex corresponds to a complex reshuffle of the mesoscopic information network"

**Giampiero Bardella[1*], Valentina Giuffrida[2], Franco Giarrocco[2], Emiliano Brunamonti[1], Pierpaolo Pani[1], and Stefano Ferraina[1]**

[1]Department of Physiology and Pharmacology, Sapienza University of Rome, Piazzale Aldo Moro 5, 00185, Rome, Italy
[2]PhD Program in Behavioral Neuroscience, Sapienza University of Rome, Piazzale Aldo Moro 5, 00185, Rome, Italy

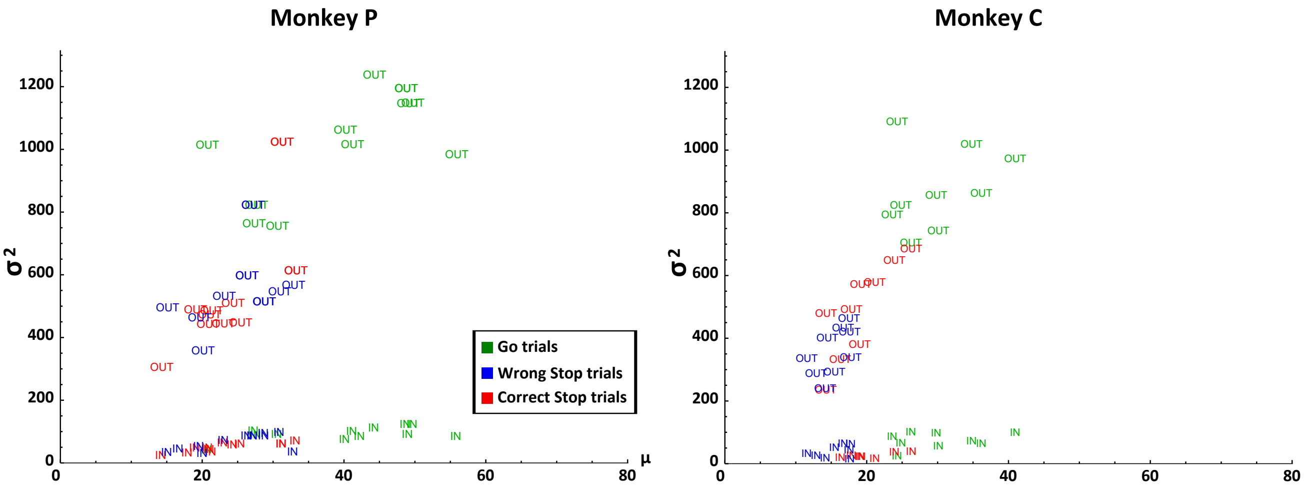


**Figure S1.** $VD_{in}$ (IN markers) and $VD_{out}$ (OUT markers) distribution parameters (x axis: mean, $\mu$; y axis: variance, $\sigma^2$) of the PMd empirical information network across behavioral conditions for each recording session for both monkeys. As evident, only the $VD_{out}$ distributions showed a great excursion of the variance with respect to the mean, i.e. show the presence of a fat-tail. Indeed, for the $VD_{out}$ distributions $\sigma^2$ is at least one order of magnitude greater than $\mu$ (Kruskal-Wallis, $p < 0.01$). This documents the presence of a fat tail and hence of high out degree nodes. On the contrary, the $VD_{in}$ distributions showed comparable values.

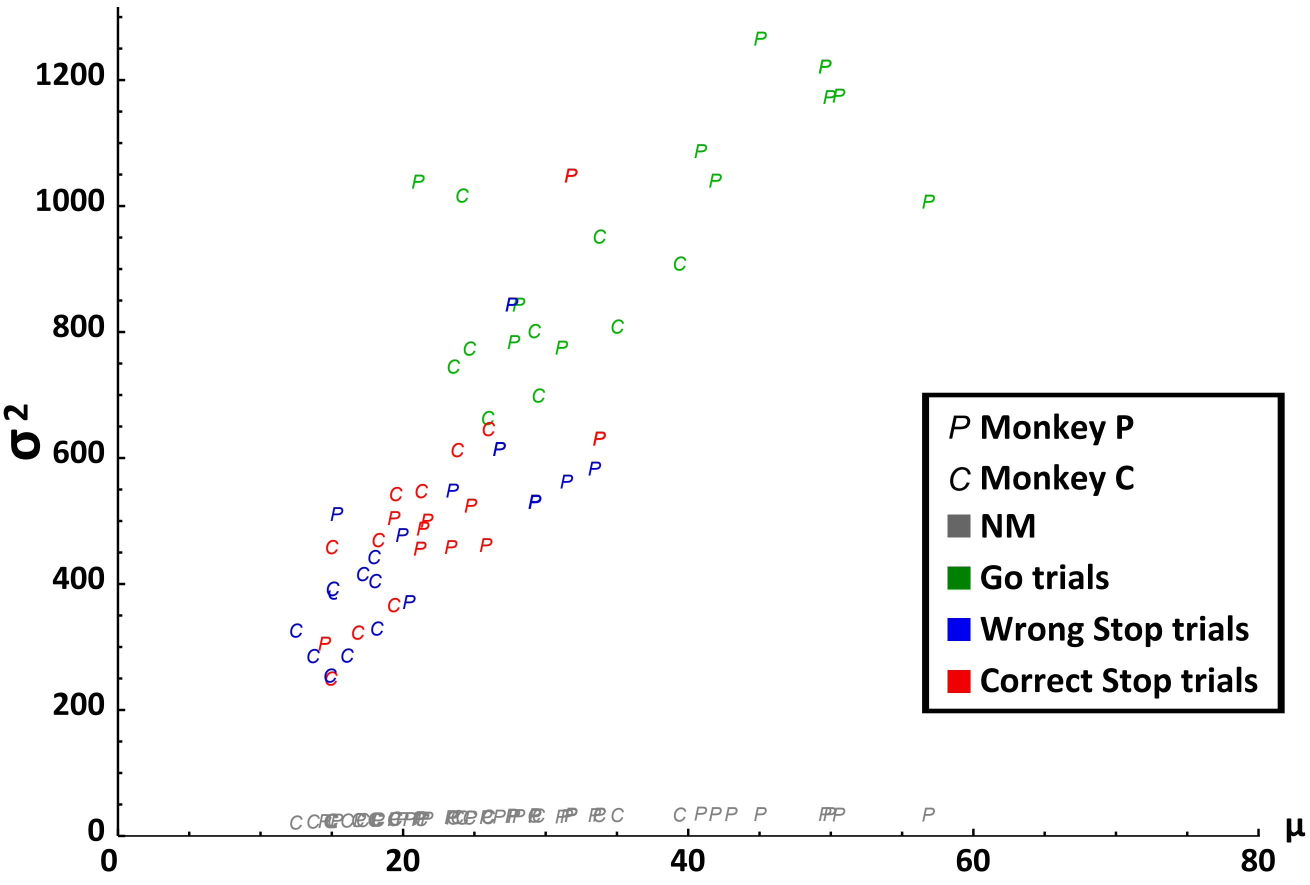


**Figure S2.** Comparison between the $VD_{out}$ distribution parameters (x axis: mean, $\mu$; y axis: variance, $\sigma^2$) of the PMd empirical information network across behavioral conditions for each recording session for both monkeys and the ensemble-average obtained from the null model (see text for details). The empirical distributions have a variance that spans values at least one order of magnitude greater than that of the networks derived from the MUA distribution, despite having comparable $\mu$ values; Kruskal-Wallis, $p < 0.01$, Bonferroni corrected for multiple comparisons. (**NM**: Null model).

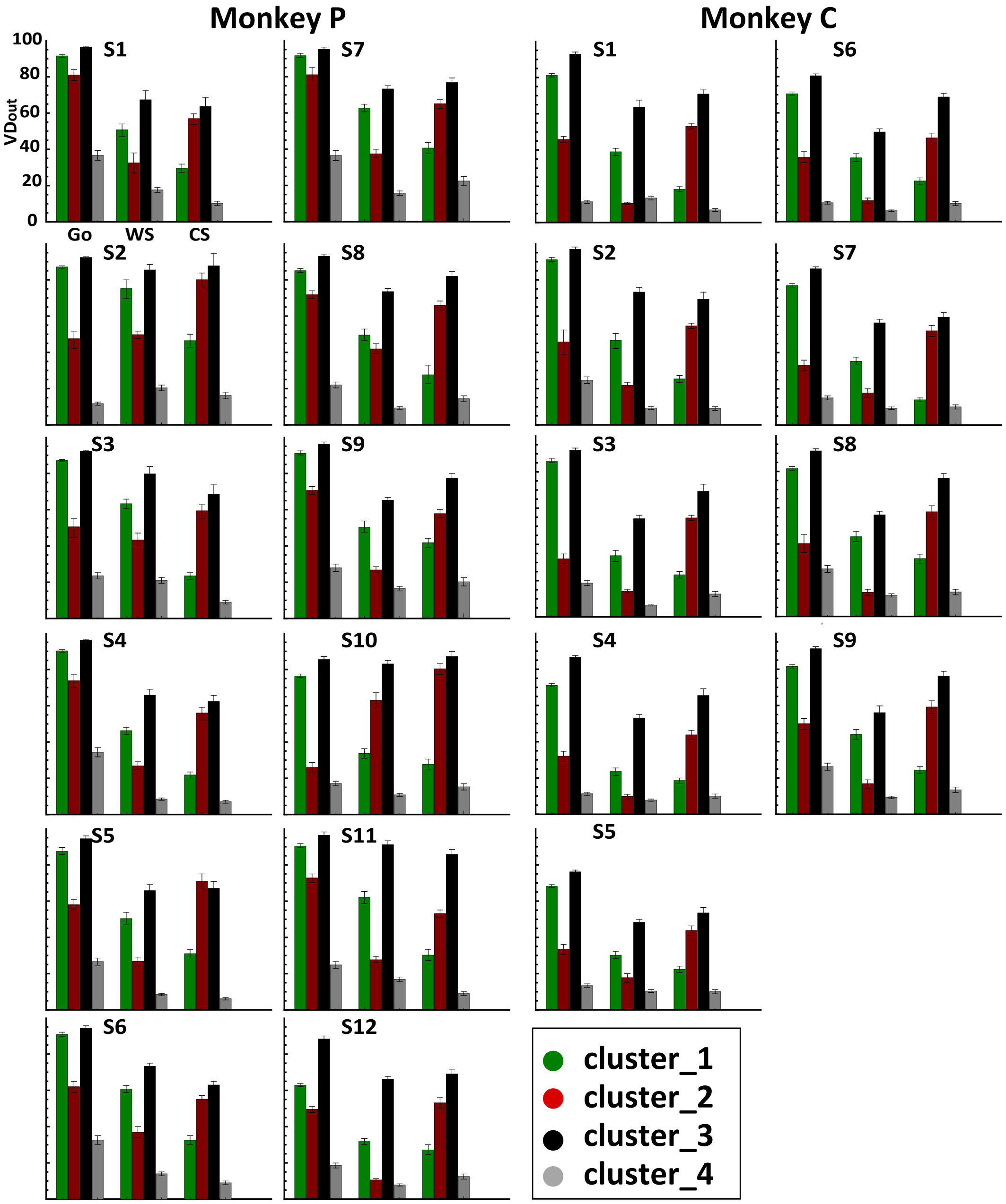


**Figure S3.** Topology of information transmission at the single-session level. $VD_{out}$ values compared between clusters and across behavioural conditions. Colours reflect the neuronal clusters (see legend). Error bar is the SEM of the nodes forming the cluster based on Kruskal-Wallis, all $p < 0.01$ Bonferroni corrected for multiple comparisons. Table S5 reports the average over sessions for each cluster. (**Go**: Go trials; **WS**: wrong Stop trials; **CS**: correct Stop trials)

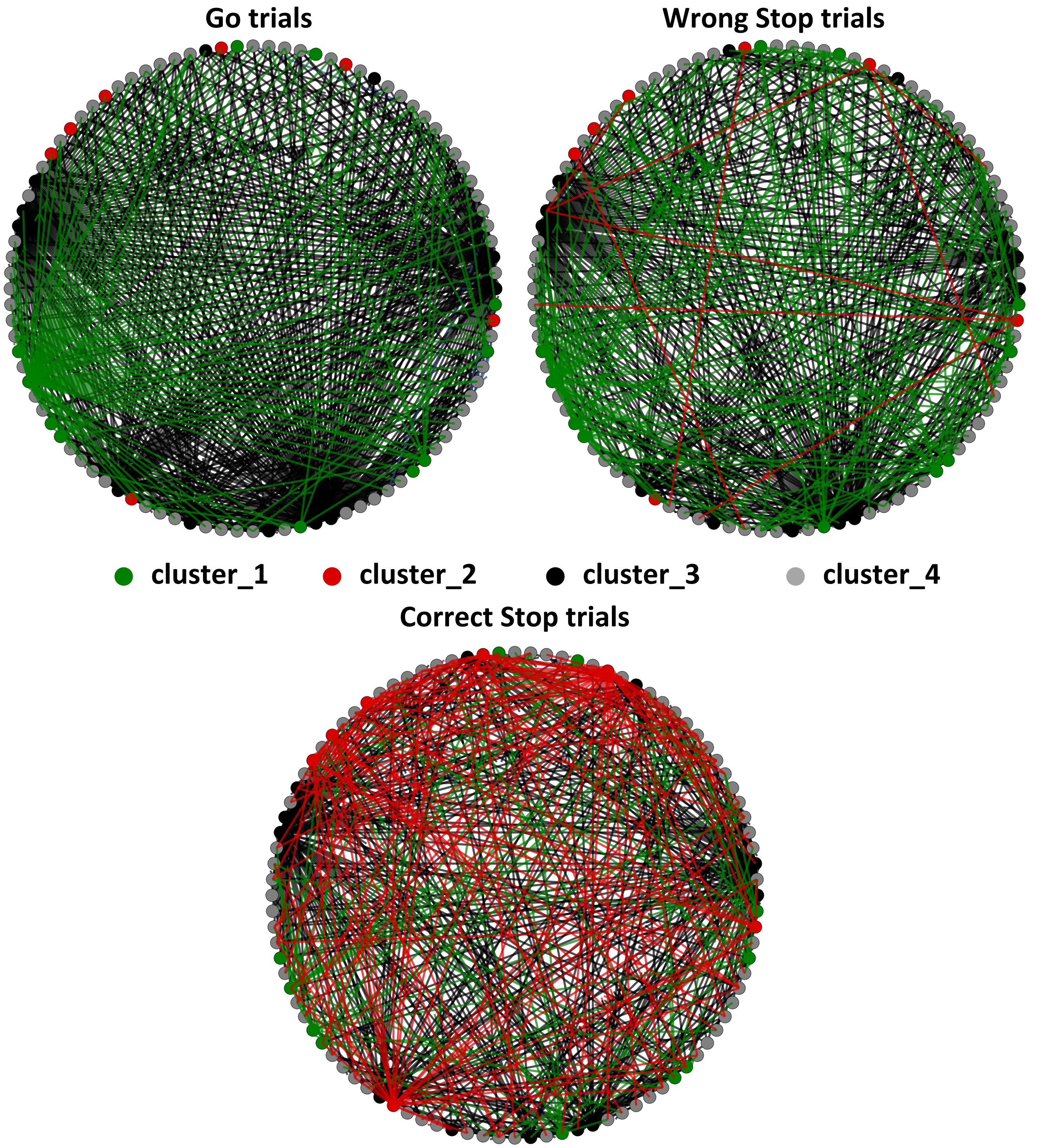


**Figure S4.** Detail of the PMd information network of Go, wrong and correct Stop trials for data in Fig.4. Each node and the respective connections are colour coded accordingly to the corresponding cluster (see legend). The coordinates of each node on the circle are the same as Fig. 4

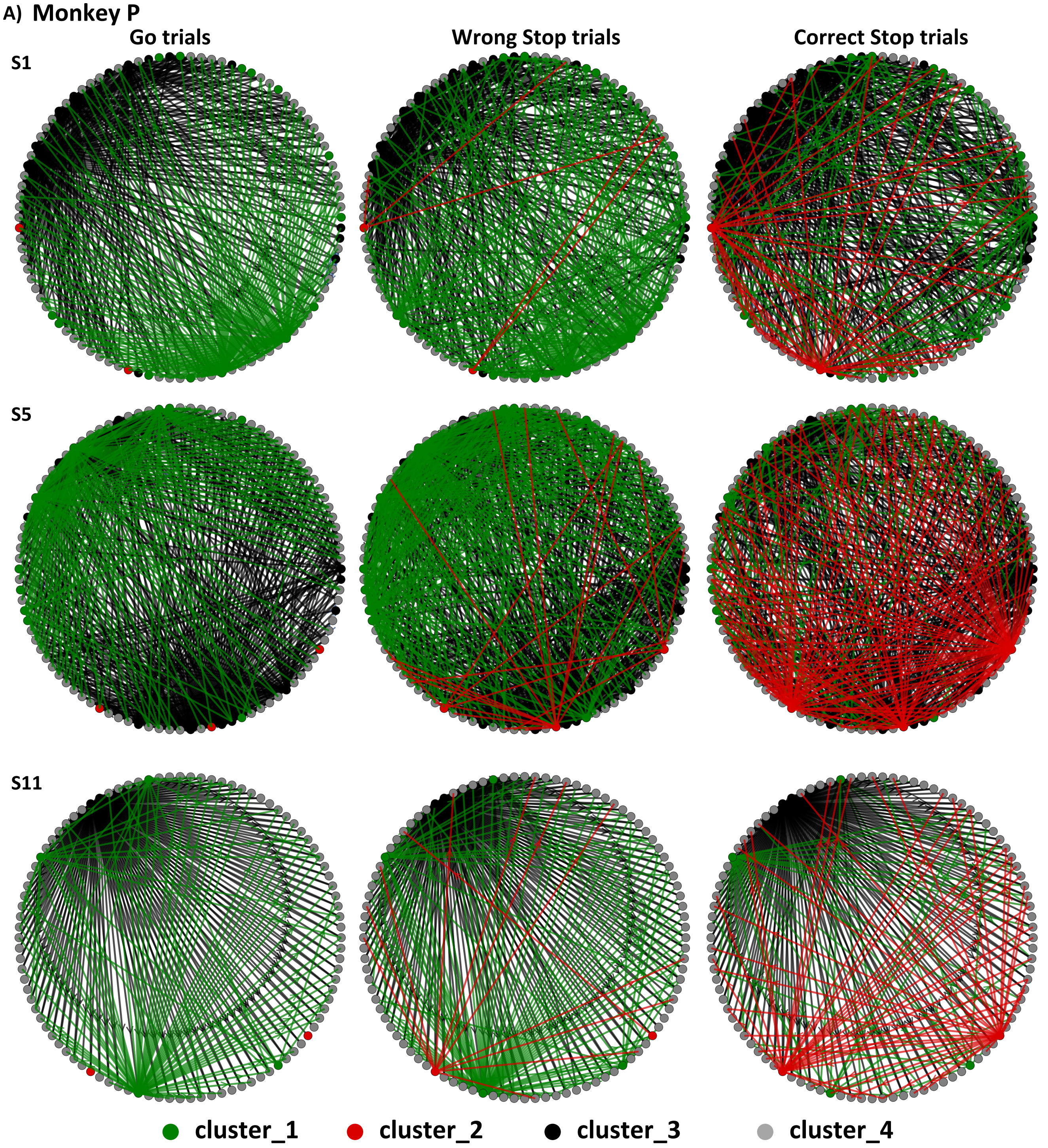

A) Monkey P
Go trials
Wrong Stop trials
Correct Stop trials
S1
S5
S11
cluster_1
cluster_2
cluster_3
cluster_4

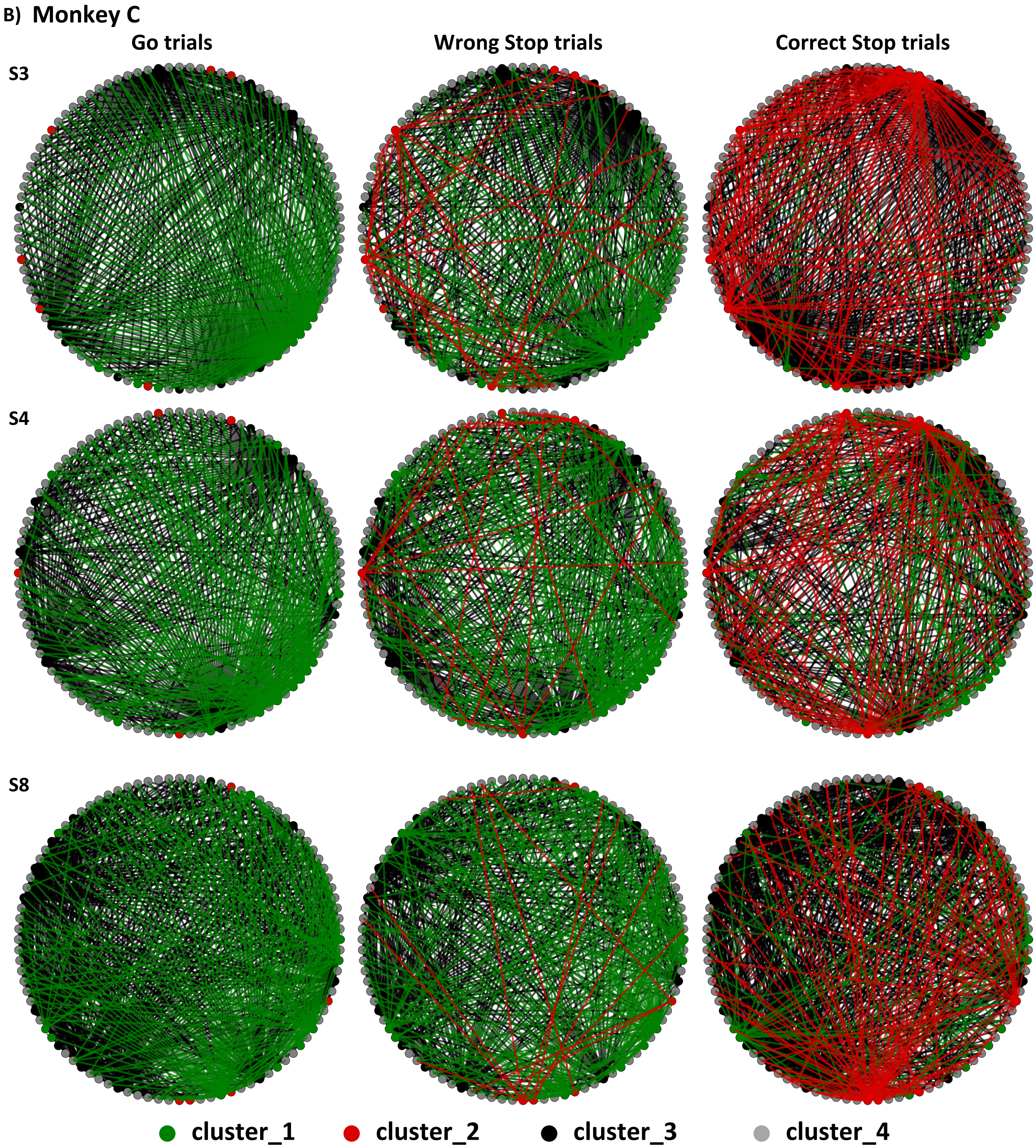


**Figure S5.** Detail of the PMd information network of Go, wrong and correct Stop trials for three example sessions for Monkey P (panel A) and Monkey C (panel B). Each node and the respective connections are colour coded accordingly to the corresponding cluster (see legend). The coordinates of each node on the circle are preserved for each network. As in Fig. 4 and Fig. S4, only the 20% of strongest connections are shown for illustrative purposes. The example sessions shown here are those that, given the network's link density, were best suited to be presented in this graphical representation. The coordinates of each node on the circle are the same as Fig. 4 and Fig. S4 and are preserved across behavioral conditions and animals.

| Behavioural Results | | | | | | | |
|---|---|---|---|---|---|---|---|
| Monkey P | | | | | | | |
| $S$ | $\overline{RT}_{Go}$ | $\overline{RT}_{Wr}$ | $\overline{SSD}_{CS}$ | $\overline{SSD}_{Wr}$ | $SSRT$ | $P_{inhibit}$ | *p-value* |
| 1 | 590 ms | 559 ms | 273 ms | 337 ms | 285 ms | 0.52 | $p < 0.05$ |
| 2 | 584 ms | 565 ms | 305 ms | 408 ms | 227 ms | 0.50 | $p < 0.05$ |
| 3 | 618 ms | 592 ms | 335 ms | 430 ms | 238 ms | 0.52 | $p < 0.05$ |
| 4 | 573 ms | 549 ms | 301 ms | 401 ms | 221 ms | 0.50 | $p < 0.01$ |
| 5 | 572 ms | 540 ms | 293 ms | 379 ms | 236 ms | 0.50 | $p < 0.05$ |
| 6 | 643 ms | 625 ms | 402 ms | 496 ms | 195 ms | 0.51 | $p < 0.05$ |
| 7 | 600 ms | 568 ms | 340 ms | 441 ms | 209 ms | 0.49 | $p < 0.01$ |
| 8 | 664 ms | 641 ms | 469 ms | 525 ms | 170 ms | 0.57 | $p < 0.01$ |
| 9 | 786 ms | 753 ms | 528 ms | 627 ms | 213 ms | 0.55 | $p < 0.01$ |
| 10 | 672 ms | 619 ms | 421 ms | 500 ms | 216 ms | 0.57 | $p < 0.01$ |
| 11 | 766 ms | 721 ms | 497 ms | 504 ms | 230 ms | 0.51 | $p < 0.01$ |
| 12 | 868 ms | 853 ms | 675 ms | 746 ms | 160 ms | 0.55 | $p < 0.05$ |
| Monkey C | | | | | | | |
| 1 | 598 ms | 523 ms | 322 ms | 383 ms | 250 ms | 0.57 | $p < 0.01$ |
| 2 | 539 ms | 460 ms | 382 ms | 378 ms | 158 ms | 0.60 | $p < 0.05$ |
| 3 | 561 ms | 522 ms | 318 ms | 403 ms | 207 ms | 0.58 | $p < 0.05$ |
| 4 | 672 ms | 631 ms | 423 ms | 507 ms | 217 ms | 0.60 | $p < 0.05$ |
| 5 | 638 ms | 612 ms | 401 ms | 512 ms | 189 ms | 0.56 | $p < 0.05$ |
| 6 | 579 ms | 533 ms | 308 ms | 426 ms | 224 ms | 0.60 | $p < 0.01$ |
| 7 | 667 ms | 620 ms | 383 ms | 498 ms | 240 ms | 0.59 | $p < 0.05$ |
| 8 | 697 ms | 672 ms | 430 ms | 546 ms | 223 ms | 0.60 | $p < 0.05$ |
| 9 | 688 ms | 662 ms | 401 ms | 547 ms | 230 ms | 0.60 | $p < 0.05$ |

**Table S1. Behavioural results**. $S$, index of the recording session. $\overline{RT}_{Go}$, mean reaction time of Go trials. $\overline{RT}_{Wr}$, mean reaction time of wrong Stop trials. $\overline{SSD}_{CS}$, mean SSD of correct Stop trials. $\overline{SSD}_{Wr}$, mean SSD of Wrong Stop trials. $SSRT$, Stop signal reaction time. $P_{inhibit}$, inhibition probability. The *p-values* result from the independence; Kolmogorov-Smirnov test, $p < 0.05$ between $RT_{Go}$ and $RT_{Wr}$ distributions. (**Go**: Go trials; **WS**: wrong Stop trials; **CS**: correct Stop trials)

| ***I* matrix details** | | | | |
|---|---|---|---|---|
| **Monkey P** | | | | |
| **Go trials** | cluster_1 | cluster_2 | cluster_3 | cluster_4 |
| cluster_1 | / | $0.08 \pm 0.03$ | $0.12 \pm 0.02$ | $1.8 \pm 0.2$ |
| cluster_2 | $0.04 \pm 0.005$ | / | $0.04 \pm 0.01$ | $0.7 \pm 0.06$ |
| cluster_3 | $0.24 \pm 0.04$ | $0.14 \pm 0.04$ | / | $4.1 \pm 0.9$ |
| cluster_4 | $0.01 \pm 0.005$ | $0.01 \pm 0.003$ | $0.02 \pm 0.007$ | / |
| **Wrong Stop trials** | cluster_1 | cluster_2 | cluster_3 | cluster_4 |
| cluster_1 | / | $0.05 \pm 0.008$ | $0.1 \pm 0.01$ | $1.6 \pm 0.3$ |
| cluster_2 | $0.05 \pm 0.008$ | / | $0.045 \pm 0.01$ | $0.7 \pm 0.1$ |
| cluster_3 | $0.25 \pm 0.04$ | $0.13 \pm 0.04$ | / | $4.0 \pm 1.2$ |
| cluster_4 | $0.02 \pm 0.004$ | $0.008 \pm 0.001$ | $0.02 \pm 0.005$ | / |
| **Correct Stop trials** | cluster_1 | cluster_2 | cluster_3 | cluster_4 |
| cluster_1 | / | $0.01 \pm 0.003$ | $0.05 \pm 0.009$ | $0.75 \pm 0.01$ |
| cluster_2 | $0.2 \pm 0.003$ | / | $0.12 \pm 0.02$ | $1.9 \pm 0.2$ |
| cluster_3 | $0.28 \pm 0.03$ | $0.10 \pm 0.02$ | / | $3.5 \pm 0.9$ |
| cluster_4 | $0.02 \pm 0.005$ | $0.004 \pm 0.001$ | $0.02 \pm 0.005$ | / |
| **Monkey C** | | | | |
| **Go trials** | cluster_1 | cluster_2 | cluster_3 | cluster_4 |
| cluster_1 | / | $0.1 \pm 0.012$ | $0.11 \pm 0.02$ | $2.2 \pm 0.45$ |
| cluster_2 | $0.016 \pm 0.004$ | / | $0.03 \pm 0.005$ | $0.53 \pm 0.06$ |
| cluster_3 | $0.25 \pm 0.06$ | $0.18 \pm 0.02$ | / | $4.2 \pm 0.9$ |
| cluster_4 | $0.004 \pm 10^{-}4$ | $0.009 \pm 0.001$ | $0.008 \pm 0.002$ | / |
| **Wrong Stop trials** | cluster_1 | cluster_2 | cluster_3 | cluster_4 |
| cluster_1 | / | $0.065 \pm 0.013$ | $0.13 \pm 0.013$ | $2.6 \pm 0.24$ |
| cluster_2 | $0.02 \pm 0.008$ | / | $0.03 \pm 0.005$ | $0.66 \pm 0.1$ |
| cluster_3 | $0.21 \pm 0.05$ | $0.18 \pm 0.02$ | / | $4 \pm 0.7$ |
| cluster_4 | $0.014 \pm 0.002$ | $0.01 \pm 0.002$ | $0.009 \pm 0.001$ | / |
| **Correct Stop trials** | cluster_1 | cluster_2 | cluster_3 | cluster_4 |
| cluster_1 | / | $0.02 \pm 0.004$ | $0.03 \pm 0.008$ | $0.72 \pm 0.1$ |
| cluster_2 | $0.13 \pm 0.015$ | / | $0.10 \pm 0.01$ | $1.9 \pm 0.2$ |
| cluster_3 | $0.3 \pm 0.03$ | $0.11 \pm 0.02$ | / | $3.8 \pm 0.5$ |
| cluster_4 | $0.018 \pm 0.002$ | $0.007 \pm 0.001$ | $0.01 \pm 0.001$ | / |

**Table S2. *I* matrix details**. Values of I for each cluster and each behavioral condition averaged (mean ± SEM) over recording sessions. The symbol / marks the absence of self loops which are excluded from the analysis (see main text).
.

| **I matrix *p-values*** | | | |
|---|---|---|---|
| **Monkey P** | | | |
| $I_{i \to j}$ | $GO-CS$ | $GO-WS$ | $CS-WS$ |
| $C_1 \to C_2$ | $p < 0.01$ | $p = 1$ | $p < 0.05$ |
| $C_1 \to C_3$ | $p < 0.01$ | $p = 1$ | $p < 0.01$ |
| $C_1 \to C_4$ | $p < 0.01$ | $p = 1$ | $p < 0.01$ |
| $C_2 \to C_1$ | $p < 0.01$ | $p = 1$ | $p < 0.05$ |
| $C_2 \to C_3$ | $p < 0.01$ | $p = 1$ | $p < 0.05$ |
| $C_2 \to C_4$ | $p < 0.01$ | $p = 1$ | $p < 0.01$ |
| $C_3 \to C_1$ | $p = 0.7$ | $p = 1$ | $p = 0.7$ |
| $C_3 \to C_2$ | $p = 0.5$ | $p = 1$ | $p = 0.8$ |
| $C_3 \to C_4$ | $p = 1$ | $p = 1$ | $p = 1$ |
| $C_4 \to C_1$ | $p = 0.2$ | $p = 1$ | $p = 0.6$ |
| $C_4 \to C_2$ | $p = 0.3$ | $p = 1$ | $p = 0.6$ |
| $C_4 \to C_3$ | $p = 1$ | $p = 1$ | $p = 15$ |
| **Monkey C** | | | |
| $C_1 \to C_2$ | $p < 0.01$ | $p = 0.7$ | $p < 0.05$ |
| $C_1 \to C_3$ | $p < 0.01$ | $p = 0.9$ | $p < 0.05$ |
| $C_1 \to C_4$ | $p < 0.01$ | $p = 0.7$ | $p < 0.01$ |
| $C_2 \to C_1$ | $p < 0.01$ | $p = 1$ | $p < 0.01$ |
| $C_2 \to C_3$ | $p < 0.05$ | $p = 1$ | $p < 0.05$ |
| $C_2 \to C_4$ | $p < 0.01$ | $p = 1$ | $p < 0.01$ |
| $C_3 \to C_1$ | $p = 0.6$ | $p = 1$ | $p = 0.3$ |
| $C_3 \to C_2$ | $p = 0.1$ | $p = 1$ | $p = 0.1$ |
| $C_3 \to C_4$ | $p = 1$ | $p = 1$ | $p = 1$ |
| $C_4 \to C_1$ | $p = 0.2$ | $p = 0.3$ | $p = 0.2$ |
| $C_4 \to C_2$ | $p = 0.8$ | $p = 1$ | $p = 0.3$ |
| $C_4 \to C_3$ | $p = 1$ | $p = 1$ | $p = 15$ |

**Table S3. *p-values* for the interactions reported in Table S2**. $I_{i \to j}$: interaction term between cluster $i$ and cluster $j$. *p-values* are obtained from Kruskal-Wallis test over recording sessions, Bonferroni corrected for multiple comparisons. ($\boldsymbol{C_1}$: cluster_1; $\boldsymbol{C_2}$: cluster_2; $\boldsymbol{C_3}$: cluster_3; $\boldsymbol{C_4}$: cluster_4; **Go**: Go trials; **WS**: wrong Stop trials; **CS**: correct Stop trials).

| ANOVA Results | | | |
|---|---|---|---|
| **Monkey P** | | | |
| $S$ | $N_{Go-CS}$ | $N_{Go-WS}$ | $N_{CS-WS}$ |
| 1 | 70% | 0 | 51% |
| 2 | 77% | 0.01% | 66% |
| 3 | 65% | 0 | 42% |
| 4 | 69% | 0 | 60% |
| 5 | 62% | 0.01% | 45% |
| 6 | 78% | 0 | 65% |
| 7 | 79% | 0.01% | 76% |
| 8 | 92% | 0 | 84% |
| 9 | 58% | 0 | 43% |
| 10 | 68% | 0 | 44% |
| 11 | 68% | 0 | 63% |
| 12 | 70% | 0 | 68% |
| **Monkey C** | | | |
| 1 | 98% | 0 | 98% |
| 2 | 92% | 0% | 92% |
| 3 | 66% | 0 | 67% |
| 4 | 67% | 0 | 67% |
| 5 | 94% | 0% | 93% |
| 6 | 92% | 0 | 92% |
| 7 | 76% | 0% | 76% |
| 8 | 85% | 0 | 84% |
| 9 | 89% | 0 | 88% |

**Table S4. ANOVA results**. $S$, index of the recording session. $N_{Go-CS}$, number of modules with $p < 0.01$ between Go and correct Stop trials. $N_{Go-WS}$, number of modules with $p < 0.01$ between Go and wrong Stop trials. $N_{CS-WS}$, number of modules with with $p < 0.01$ between correct and wrong Stop trials. All $Ns$ are in percentage over the total number of moduels. *p-values* are Bonferroni corrected for multiple comparisons. (**Go**: Go trials; **WS**: wrong Stop trials; **CS**: correct Stop trials)

| Clusters Composition | |
|---|---|
| **Monkey P** $N = 96$ | |
| **Cluster** | $\mu \pm SEM$ |
| cluster_1 | $9.3 \pm 1.06$ |
| cluster_2 | $3.6 \pm 1.4$ |
| cluster_3 | $11.8 \pm 1.6$ |
| cluster_4 | $71.3 \pm 2.3$ |
| **Monkey C** $N = 96$ | |
| **Cluster** | $\mu \pm SEM$ |
| cluster_1 | $7.4 \pm 0.8$ |
| cluster_2 | $4.1 \pm 0.6$ |
| cluster_3 | $9.4 \pm 1.1$ |
| cluster_4 | $75.0 \pm 1.8$ |

**Table S5. Clusters composition**. For each monkey the composition of clusters averaged (mean ± SEM) over recording sessions is reported. $N$ is the number of modules available.

| $VD_{out}$ | | | |
|---|---|---|---|
| **Monkey P** $N = 12$ | | | |
| **Cluster** | **Go trials** | **Wrong Stop trials** | **Correct Stop trials** |
| cluster_1 | $87.74 \pm 0.77$ | $52 \pm 1.75$ | $32.86 \pm 1.33$ |
| cluster_2 | $48.74 \pm 3.5$ | $43.23 \pm 3.11$ | $67.14 \pm 2.43$ |
| cluster_3 | $92.02 \pm 0.46$ | $68.6 \pm 1.54$ | $72.77 \pm 1.34$ |
| cluster_4 | $26.04 \pm 0.81$ | $14.31 \pm 0.40$ | $13.96 \pm 0.39$ |
| **Monkey C** $N = 9$ | | | |
| **Cluster** | **Go trials** | **Wrong Stop trials** | **Correct Stop trials** |
| cluster_1 | $77.73 \pm 1.49$ | $35.56 \pm 1.80$ | $24.44 \pm 1.33$ |
| cluster_2 | $37.78 \pm 2.09$ | $17.27 \pm 1.32$ | $52.08 \pm 1.84$ |
| cluster_3 | $84.82 \pm 1.27$ | $54.95 \pm 2.50$ | $69.62 \pm 1.92$ |
| cluster_4 | $17.06 \pm 0.59$ | $8.37 \pm 0.24$ | $10.28 \pm 0.31$ |

**Table S6** Details of $VD_{out}$ across behavioral conditions. For each monkey $VD_{out}$ values averaged (mean ± SEM) over recording sessions are reported. The cluster_3 showed the highest values of $VD_{out}$ compared to other clusters in all behavioural conditions. The cluster_1 and the cluster_2 showed the second highest $VD_{out}$ values during both Go and wrong Stop trials and correct Stop trials respectively; Kruskal-Wallis, $p < 0.01$ Bonferroni corrected for multiple comparisons. **N**: number of session available.
